# Targeting Reductive Metabolic Shifts by T315I Mutation in BCR-ABL Myeloid Leukemia for Therapy

**DOI:** 10.1101/2022.03.22.485260

**Authors:** Chang-Yu Huang, Yin-Hsuan Chung, Sheng-Yang Wu, Hsin-Yang Wang, Zhi-Yu Lin, Tsung-Jung Yang, Jim-Ming Feng, Chun-Mei Hu, Zee-Fen Chang

## Abstract

T315I mutation of Bcr-Abl in chronic myeloid leukemia (CML) leads to therapeutic resistance. It is known that Bcr-Abl transformation causes ROS-induced DNA damages and replication stress, which can be exploited for anti-nucleotide therapy. We developed a small compound, JMF4073, which inhibited pyrimidylate kinases and selectively eliminated Bcr-Abl-transformed, but not untransformed myeloid cells, due to dTTP exhaustion and ROS-induced replication stress. However, T315I-Bcr-Abl-transformed cells were less vulnerable to JMF4073 because of higher dTTP pool and low replication stress. Unlike WT-Bcr-Abl-transformed cells, T315I-Bcr-Abl cells lacked Sirt1- regulated OXPHOS with increased glutamine flux to reductive carboxylation in TCA cycle and glutathione synthesis. Blocking mitochondrial pyruvate carrier (MPC) by UK-5099 reduced NADH and glutathione levels with replication stress induction, thereby converting T315I-Bcr-Abl cells sensitive to JMF4073 with dTTP and dCTP depletion. The combination of JMF4073 with UK-5099 showed in vivo eradication of T315I-Bcr-Abl-CML. These data reveal that T315I mutation causes reductive metabolic shifts in Bcr-Abl-CML, and demonstrate the therapeutic option by co-targeting MPC and pyrimidylate kinases.

## Introduction

The t(9;22) chromosomal reciprocal translocation, also known as Philadelphia chromosome (Ph), generates *BCR-ABL* fusion gene to cause chronic myeloid leukemia (CML) (Rowley, 1973, Sawyers, 1999). *BCR-ABL* encodes a constitutively active tyrosine kinase that phosphorylates a panel of downstream targets to cause uncontrolled cell proliferation, protein synthesis, and anti-apoptotic signals (Dorsey, Cunnick et al., 2002, Ren, 2005, Salesse & Verfaillie, 2002, Sattler, Mohi et al., 2002, Xie, Wang et al., 2001). Although Bcr-Abl inhibitors such as imatinib, nilotinib and dasatinib are effective in CML therapy (Braun, Eide et al., 2020, Deguchi, Kimura et al., 2008, Druker, Talpaz et al., 2001, Druker, Tamura et al., 1996), the acquisition of T315I mutation of *BCR-ABL* in patients causes the resistance to most of tyrosine kinase inhibitors (TKI), leading to poor outcome. A structure-based designed inhibitor of Bcr-Abl-T315I, ponatinib, was developed and used in the clinical trial. However, the side effects were severe (Braun et al., 2020, Leong, Aghel et al., 2021, Rosti, Castagnetti et al., 2017). While the biochemical analyses reveal the tyrosine kinase activity of T315I-Bcr-Abl lower than that of wild-type-Bcr-Abl (Skaggs, Gorre et al., 2006), a variety of studies have shown that T315I might enhance or reduce the kinase activity in vivo depending on the model used for the investigation and confer the substrate differences of Bcr-Abl in transformation (Griswold, MacPartlin et al., 2006, Mian, Schull et al., 2009). Presumably, these molecular differences change the cellular behaviors of transformed cells. This study examined the metabolic changes by T315I mutation in Bcr-Abl-transformed cells, aiming to develop anti-metabolic therapy for eradicating T315I-Bcr-Abl leukemia.

It has been shown that Bcr-Abl-transformation in myeloid progenitor cells elevates reactive oxygen species (ROS) to induce DNA double-strand breaks (DSBs), thereby leading to genome instability (Nowicki, Falinski et al., 2004). Replication stress can increase DNA double-strand breaks and facilitate the genome evolution in cancer (Brown, Spinelli et al., 2017, D’Angiolella, Donato et al., 2012, Hu, Tsao et al., 2019, Le, Poddar et al., 2017, Lecona & Fernandez-Capetillo, 2018, Niida, Katsuno et al., 2010). Mitochondrial dysfunction is the major source in ROS overproduction, and a number of oncogenic signals have been found to cause mitochondrial dysfunction (Hu, Lu et al., 2012b, Sabharwal & Schumacker, 2014, Ward & Thompson, 2012). It has been reported that Bcr-Abl promotes Rac2-GTPase activity to affect the flux of electron transfer chain complex III in mitochondria, in turn elevating ROS in Bcr-Abl cells (Nieborowska-Skorska, Kopinski et al., 2012). However, on the other hand, upregulation of OXPHOS activation in mitochondria has been shown critical for CML progression and TKI resistance (Abraham, Qiu et al., 2019, Alvarez-Calderon, Gregory et al., 2015, de Beauchamp, Himonas et al., 2022, Flis, Irvine et al., 2012, Kuntz, Baquero et al., 2017). Up-to-date, it remains unclear whether T315I mutation would change BCR-Abl-induced genome stress, mitochondrial regulation, or metabolic rewiring in myeloid cells. Understanding the alterations caused by T315I mutation would inform the therapeutic opportunity for CML acquiring T315I mutation.

A recent study has reported that the increase of OXPHOS function confers cancer cells pyrimidine biosynthesis through dihydroorotate dehydrogenase (DHODH) associated with electron transfer chain in mitochondria for proliferation (Bajzikova, Kovarova et al., 2019). Together with the one carbon folate pathway in mitochondria (Locasale, 2013, Yang & Vousden, 2016), it is possible that upregulation of OXPHOS activity in Bcr-Abl-transformed cells might benefit the dNTPs demand for uncontrolled proliferation. However, uncontrolled proliferation associated with oncogene overexpression often exhausts dNTP pools, leading to replication fork stalling, double-strand breaks, and genome instability (Halazonetis, Gorgoulis et al., 2008, Hills & Diffley, 2014). Compelling evidence has established that replication stress and DNA damage upregulate dNTP synthesis via different signal pathways to overcome replication stress and facilitate DNA repair (Brown et al., 2017, D’Angiolella et al., 2012, Le et al., 2017, Lecona & Fernandez-Capetillo, 2018). In accordance, targeting the protection signal or nucleotide synthesis has been exploited to improve anti-cancer treatment. Here, we asked whether ROS-induced replication stress and DNA damages harness Bcr-Abl-transformed cells vulnerable to dTTP inhibition, and whether T315I- Bcr-Abl transformed cells are also under replication stress in CML for metabolic targeting.

Among four dNTP, the pool size of dTTP is very the largest (Zhang, Tan et al., 2011), because high level of dTTP limits dUTP misincorporation into DNA (Chen, Tsao et al., 2016a, Hu, Yeh et al., 2012a, Vertessy & Toth, 2009). We have previously identified a compound YMU1, an inhibitor of thymidylate kinase, which increased doxorubicin sensitivity of cancer cell by causing DNA repair toxicity (Chen, Hsu et al., 2016b, Hu et al., 2012a). Using YMU1 as a lead compound, we further developed JMF4073 that gave better solubility and potency. JMF4073 inhibits both thymidylate and cytidylate kinase, which are responsible for converting dTMP and dCMP, respectively, to dTDP and dCDP for dTTP and dCTP formation in DNA replication. In this study, our data reveal the differential responses to JMF4073 in wild-type and T315I Bcr-Abl-transformed cells due to the differences in redox state, dNTP deficiency and replication stress. T315I mutation in Bcr-Abl-transformed cells causes the loss of Sirt1- regulated OXPHOS with a reductive metabolic shift. Importantly, blocking mitochondrial pyruvate carrier by UK-5099 converted these cells to acquire ROS-induced replication stress and to become sensitive to JMF4073. Finally, we showed that the combination of UK-5099 with JMF4073 prolonged the survival of T315I-Bcr-Abl CML mice.

## Results

### JMF4073 is an inhibitor of TMPK and CMPK

The survival and growth of cancer cells require sufficient supply of four dNTPs for DNA repair and replication. The reaction catalyzed by thymidylate kinase (TMPK) is essential for both de novo and salvage synthesis of TTP, while cytidylate kinase (CMPK) is essential for salvage synthesis of dCTP (Fig 1A). We have previously identified a TMPK inhibitor, namely YMU1. Using YMU1 as a lead, the compounds JMF2977 and JMF4073 were synthesized. IC_50_ of YMU1, JMF2977, and JMF4073 were 0.8, 0.5, and 0.16 μM, respectively. JMF4073 containing a fluoride substitution at C position of the pyridine ring has the lowest IC_50_ value and higher solubility as indicated clog*P* value (Fig 1B). By molecular simulation, we have previously shown that YMU1 acts in the catalytic pocket of TMPK that expel the lid domain, resulting in an open conformation that inhibits the catalytic process (Chen et al., 2016b). Since CMPK is another pyrimidylate kinase that also has a lid domain (Segura-Pena, Sekulic et al., 2004), we then tested whether JMF4073 is an inhibitor of CMPK. The results showed that JMF4073 inhibits both TMPK and CMPK at a similar IC50, 0.17 μM, whereas YMU1 is less potent to CMPK (Fig. 1B). Thus, JMF4073 is indeed an inhibitor of pyrimidylate kinases. We then determined the mode of JMF4073 inhibition by pre-incubating purified TMPK or CMPK protein with various concentrations of JMF4073, and measured the alteration in Vmax and *Km* by NADH-coupled enzymatic assay. The kinetic data demonstrated that JMF4073 inhibits TMPK and CMPK in a mixed inhibitory mode as indicated by the increased *Km* for the substrates with decreased Vmax. The inhibition constant (Ki) of JMF4073 in the dTMP and ATP kinetic analysis of TMPK was 0.16 μM and 0.005 μM, respectively. The Ki in the CMP and ATP kinetic analysis of CMPK was 0.37 μM and 0.06 μM, respectively (Table 1). These kinetic analyses indicate that JMF4073 has a higher affinity to the ATP binding sites in TMPK and CMPK.

**Figure 1.**
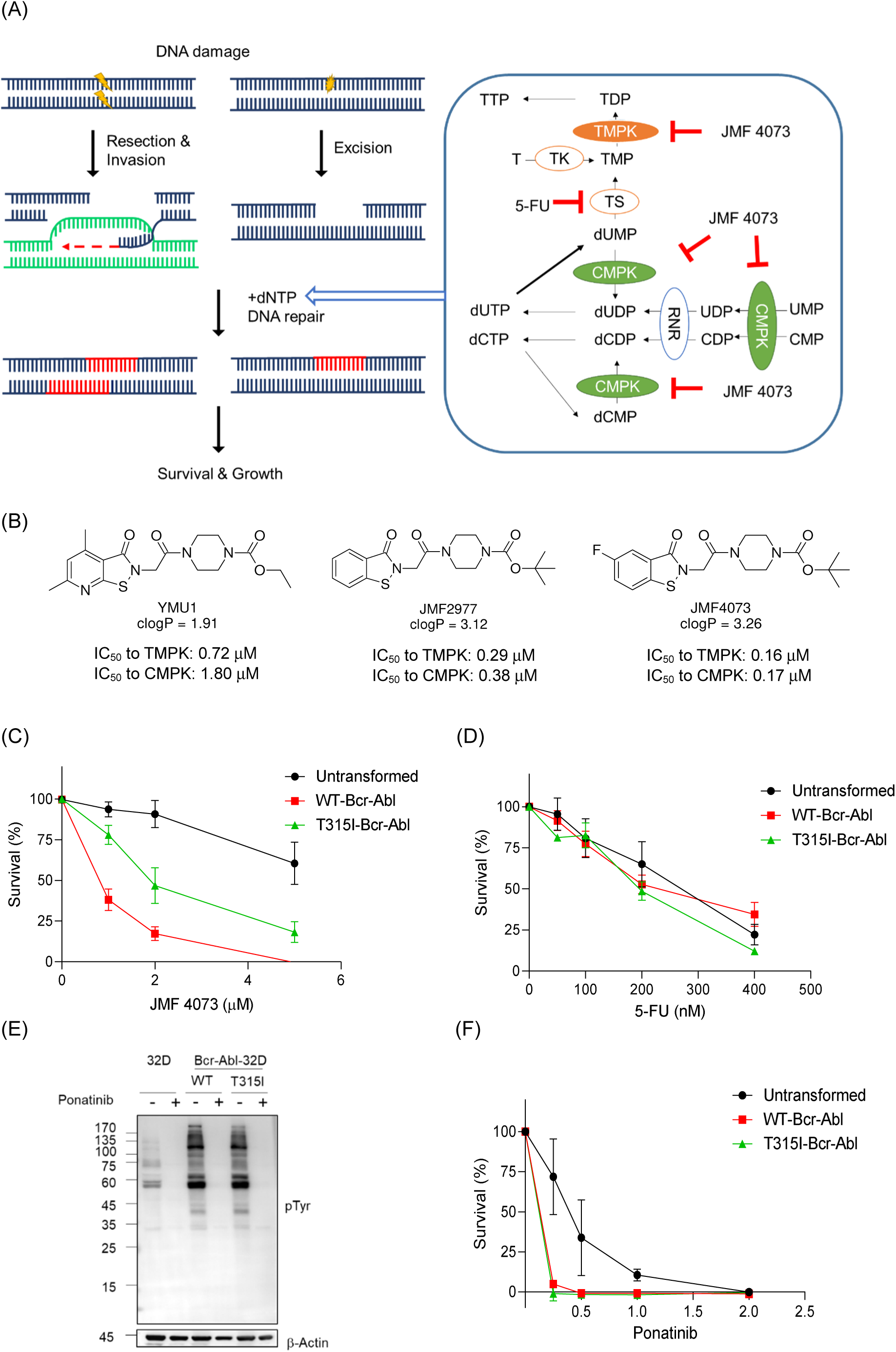
Selective susceptibility of WT- Bcr-Abl-transformed cells to JMF4073. (A)Schematic view of the requirement of dNTP supply for the survival of cells with DNA damages. TMPK and CMPK mediate the de novo and salvage synthesis of dTDP and dCDP, respectively, which subsequently are converted to dTTP and dCTP by nucleoside diphosphate kinase (NDPK). Thymidylate synthase (TS) converts dUMP to dTMP, which is responsible for the de novo synthesis of dTTP. Ribonucleotide reductase (RNR) mediates the de novo synthesis of dCDP, dADP and dGDP. Salvage synthesis of dTMP and dCMP is mediated by thymidine kinase and deoxycytidine kinase. (B) The chemical structures, clogP and IC50 of YMU1, JMF 2977, and JMF4073. The clogP values were calculated by Chemsketch. The IC50 values against hTMPK and hCMPK were measured by NADH-coupled TMPK assay. The results are shown as averages ± SEM (n = 3). (C-D) WT-, and T315I-Bcr-Abl-transformed and unstransformed 32D myeloid progenitor cells were treated with (C) JMF4073 or (D) 5- FU at the indicated concentrations. After three days, viability assays by CCK-8 were performed. Data are represented as mean ± SEM from three independent experiments. (E) The comparison of the overall pTyr in unstranformed, WT, and T315I Bcr-Abl - transformed 32D myeloid progenitor cells after the treatment of ponatinib (2 μM) for 4h (F) The cell viability was measured after 3 days of ponatinib treatment at the indicated concentration. Data are represented as mean ± SEM from three independent experiments.

**Table 1.**
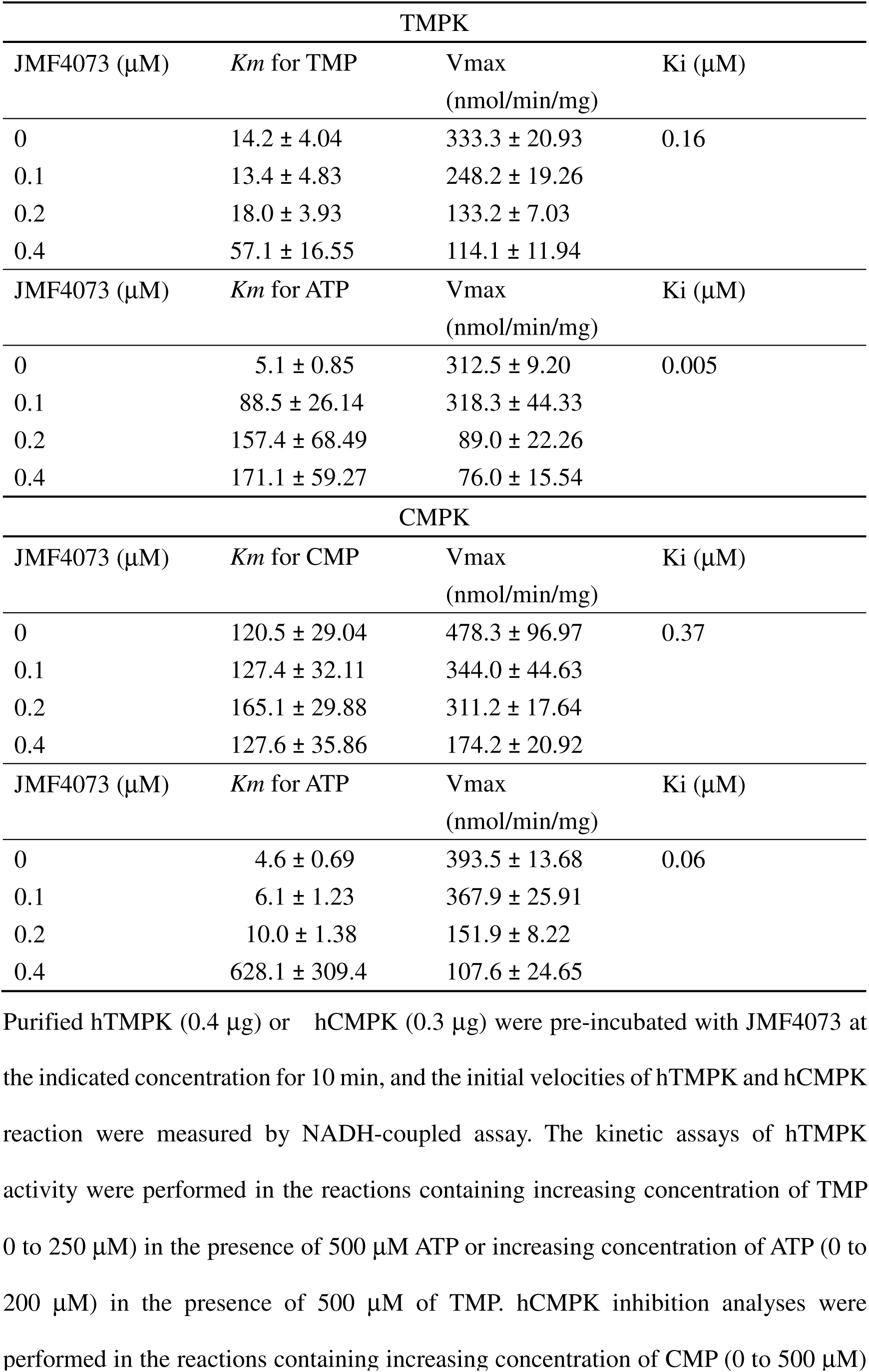
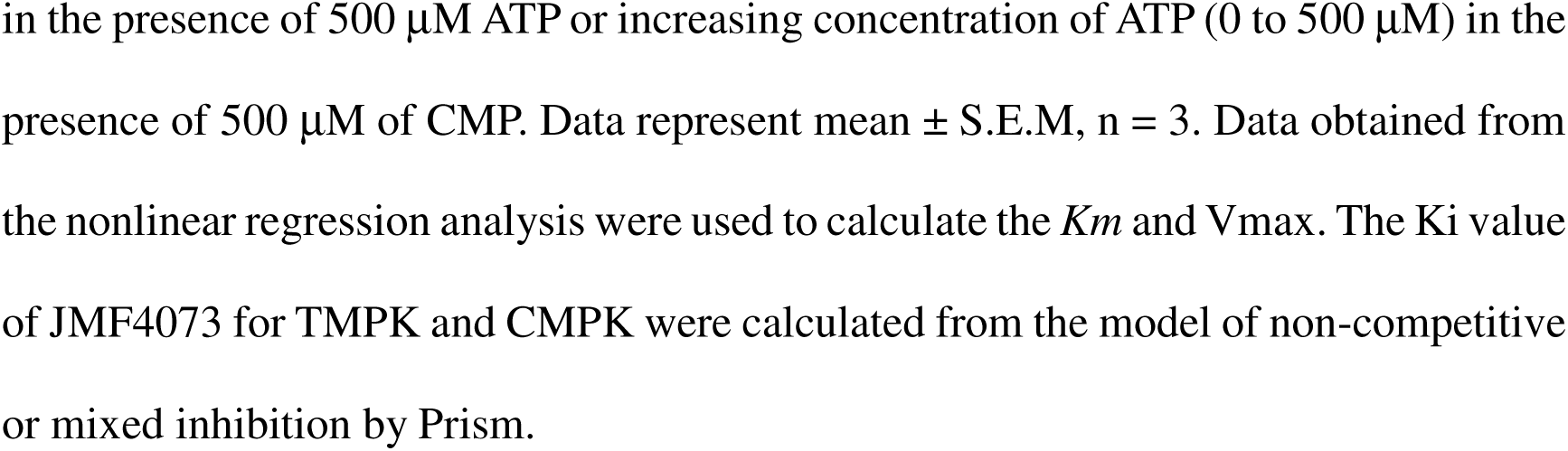
Effects of JMF4073 on enzymatic kinetics of purified human TMPK and CMPK.

| TMPK |  |  |  |
| --- | --- | --- | --- |
| JMF4073 ($\mu\text{M}$) | $K_m$ for TMP | $V_{\text{max}}$<br>(nmol/min/mg) | $K_i$ ($\mu\text{M}$) |
| 0 | $14.2 \pm 4.04$ | $333.3 \pm 20.93$ | 0.16 |
| 0.1 | $13.4 \pm 4.83$ | $248.2 \pm 19.26$ | |
| 0.2 | $18.0 \pm 3.93$ | $133.2 \pm 7.03$ | |
| 0.4 | $57.1 \pm 16.55$ | $114.1 \pm 11.94$ | |
| JMF4073 ($\mu\text{M}$) | $K_m$ for ATP | $V_{\text{max}}$<br>(nmol/min/mg) | $K_i$ ($\mu\text{M}$) |
| 0 | $5.1 \pm 0.85$ | $312.5 \pm 9.20$ | 0.005 |
| 0.1 | $88.5 \pm 26.14$ | $318.3 \pm 44.33$ | |
| 0.2 | $157.4 \pm 68.49$ | $89.0 \pm 22.26$ | |
| 0.4 | $171.1 \pm 59.27$ | $76.0 \pm 15.54$ | |
| CMPK |  |  |  |
| JMF4073 ($\mu\text{M}$) | $K_m$ for CMP | $V_{\text{max}}$<br>(nmol/min/mg) | $K_i$ ($\mu\text{M}$) |
| 0 | $120.5 \pm 29.04$ | $478.3 \pm 96.97$ | 0.37 |
| 0.1 | $127.4 \pm 32.11$ | $344.0 \pm 44.63$ | |
| 0.2 | $165.1 \pm 29.88$ | $311.2 \pm 17.64$ | |
| 0.4 | $127.6 \pm 35.86$ | $174.2 \pm 20.92$ | |
| JMF4073 ($\mu\text{M}$) | $K_m$ for ATP | $V_{\text{max}}$<br>(nmol/min/mg) | $K_i$ ($\mu\text{M}$) |
| 0 | $4.6 \pm 0.69$ | $393.5 \pm 13.68$ | 0.06 |
| 0.1 | $6.1 \pm 1.23$ | $367.9 \pm 25.91$ | |
| 0.2 | $10.0 \pm 1.38$ | $151.9 \pm 8.22$ | |
| 0.4 | $628.1 \pm 309.4$ | $107.6 \pm 24.65$ | |

Purified hTMPK (0.4 $\mu\text{g}$ ) or hCMPK (0.3 $\mu\text{g}$ ) were pre-incubated with JMF4073 at the indicated concentration for 10 min, and the initial velocities of hTMPK and hCMPK reaction were measured by NADH-coupled assay. The kinetic assays of hTMPK activity were performed in the reactions containing increasing concentration of TMP (0 to 250 $\mu\text{M}$ ) in the presence of 500 $\mu\text{M}$ ATP or increasing concentration of ATP (0 to 200 $\mu\text{M}$ ) in the presence of 500 $\mu\text{M}$ of TMP. hCMPK inhibition analyses were performed in the reactions containing increasing concentration of CMP (0 to 500 $\mu\text{M}$ )
in the presence of 500 $\mu\text{M}$ ATP or increasing concentration of ATP (0 to 500 $\mu\text{M}$ ) in the presence of 500 $\mu\text{M}$ of CMP. Data represent mean ± S.E.M, $n = 3$ . Data obtained from the nonlinear regression analysis were used to calculate the $K_m$ and $V_{\text{max}}$ . The $K_i$ value of JMF4073 for TMPK and CMPK were calculated from the model of non-competitive or mixed inhibition by Prism.

### Different JMF4073 susceptibility in WT- and T315I-Bcr-Abl-transformed and untransformed myeloid cells

The supply of dNTP is critical for overcoming DNA damages and replication stress for survival. We then tested the effect of JMF4073 on the growth of WT-, T315I-Bcr-Abl-transformed and untransformed 32D cells. The results showed that WT-Bcr-Abl cells were highly susceptible to JMF4073 with GI_50_ 0.67 μM as compared to 2.0 μM and 6.81 μM for T315I-Bcr-Abl- and untransformed 32D cells, respectively (Fig 1C). Overexpression of TMPK and CMPK by lentiviral infection in Bcr-Abl-32D cells reduced JMF4073 sensitivity, indicating the involvement of inhibition of TMPK and CMPK in the susceptibility to JMF4073 (Appendix Fig S1). 5-FU is a widely used anti-nucleotide drug that targets thymidylate synthase to induce dUTP toxicity. 5-FU sensitivity was similar in WT-, T315I-Bcr-Abl-32D and untransformed 32D progenitor cells (Fig. 1D). Since inhibition of TMPK or CMPK would not generate dUTP toxicity, 5-FU is clearly more toxic than JMF4073 to untransformed 32D cells. Treatment with ponatinib, which is an inhibitor of WT- and T315I-Bcr-Abl, decreased overall tyrosine phosphorylation of cellular proteins and cell viability not only in WT- and T315I-Bcr-Abl-transformed but also 32D progenitor cells (Fig 1E-1F), suggesting the general toxicity. Thus, JMF4073 treatment has an advantage over 5-FU and ponatinib in selectively suppressing Bcr-Abl-transformed but not untransformed myeloid progenitor cells.

Since constitutive tyrosine kinase activation is responsible the uncontrolled growth of CML, the effect of JMF4073 on global tyrosine phosphorylation was examined. JMF4073 treatment for 16 h did not reduce overall tyrosine phosphorylation, but treatment for 6 h was sufficient to reduce dTTP and dCTP pools (Fig 2A-2B). Lesser effect on reducing dTTP and dCTP pool was observed in T315I-Bcr-Abl cells. As mentioned earlier, JMF4073 has a high affinity to ATP binding site of CMPK and TMPK. To test whether cellular level of ATP might contribute to JMF4073 effect, we measured ATP amount in these cells. It appeared that T315I-Bcr-Abl cells contained 3- fold higher level of ATP than WT-Bcr-Abl- and untransformed 32D cells (Fig 2C). It is possible that high ATP level in T315I-Bcr-Abl cells affect the efficacy of JMF4073 on inhibition of CMPK and TMPK. We further tested the in vivo therapeutic effect of JMF4073 in WT- and T315I-Bcr-Abl-CML mice. WT-Bcr-Abl-transformed 32D cells were transplanted to C3H/HeNCrNarl mice. After 48 h, these mice were treated with a daily intraperitoneal injection of JMF4073 for 14 days (Fig 2D). After the therapy, the hematology analysis showed white blood counts were brought down by JMF4073 treatment in WT-Bcr-Abl-32D bearing mice (Appendix Table S1). Moreover, the 14- day-JMF4073 treatments prolonged the survival of leukemia mice (Fig 2D). Since T315I-Bcr-Abl-transformed 32D cells took longer time to develop leukemia phenotype than WT Bcr-Abl cells, we transplanted 2-fold more cells into C3H/HeNCrNarl mice. After transplantation for 7 days, these mice were treated with JMF4073 for 14 days. Unlike WT-Bcr-Abl mice, the survival of T315I leukemia mice was not significantly prolonged by JMF4073 (Fig 2E). Thus, unlike WT-Bcr-Abl cells, T315I-Bcr-Abl cells are less vulnerable to JMF4073 in vitro and in vivo.

**Figure 2.**
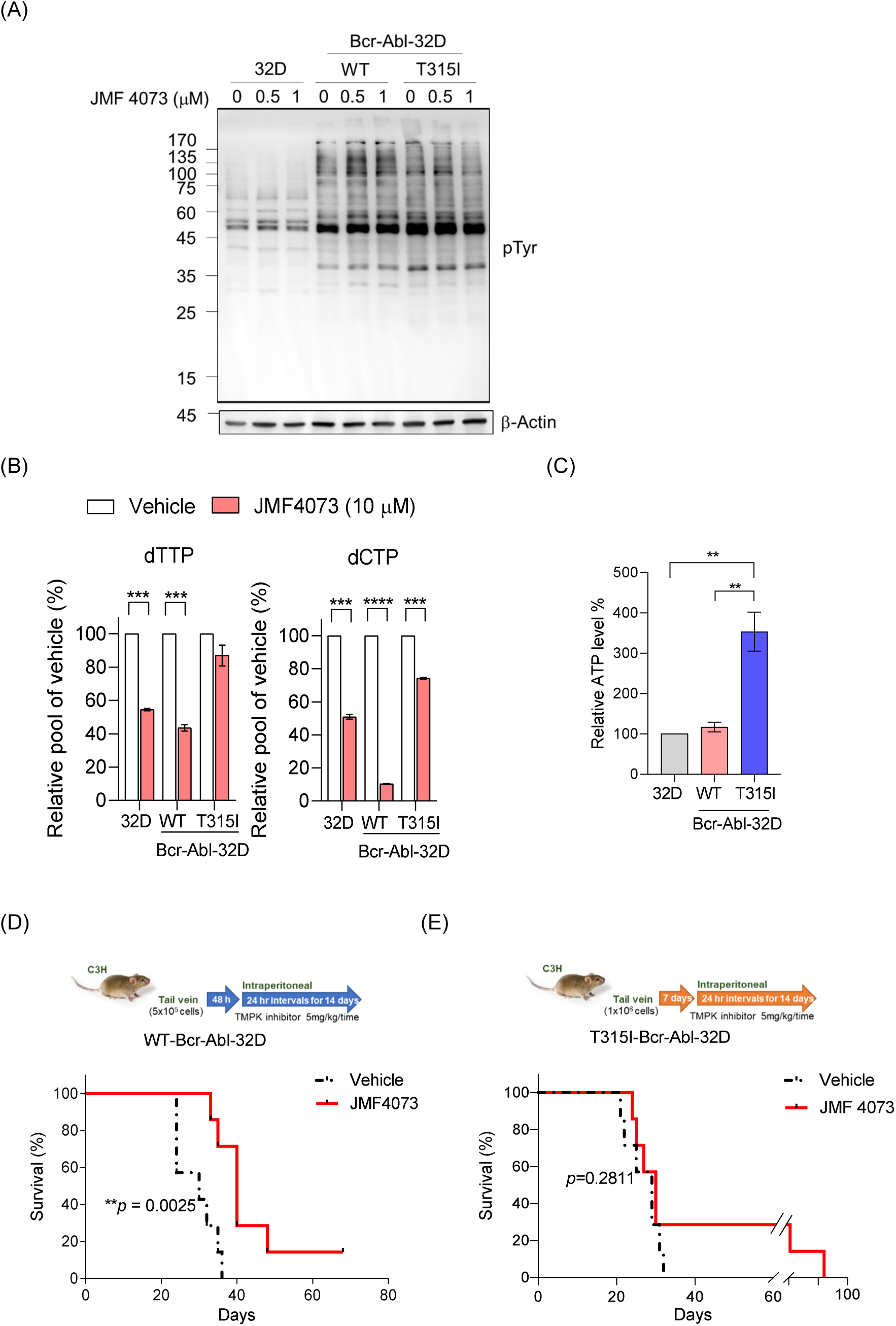
T315I mutation impacts susceptibility to JMF4073 in vitro and in vivo. (A) The comparison of the overall tyrosine phosphorylation in untransformed, WT-, and T315I-Bcr-Abl transformed 32D cells after treatment with JMF4073 for 16h. (B) The alteration of dTTP and dCTP pools in untransformed, WT-, and T315I-Bcr-Abl transformed 32D cells after incubation with JMF4073 (10 μM) for 6h. Data are represented as mean ± SEM from three independent experiments. (C) Untransformed, WT-, T315I-Bcr-Abl-32D cells were harvested for ATP measurement. Data are presented as mean ± SEM from at least three independent experiments. Asterisks denote *p < 0.05, **p< 0.01 by non-paired t-test. (D) C3H/HeNCrNarl mice were intravenously injected with 5 × 10^5^ cells of WT-Bcr-Abl-32D cell. After 48 hr of transplantation, mice were treated with vehicle (n = 7), or JMF4073 (5 mg/kg/time, n = 7) by intraperitoneal injection at 24 h interval for 14 days. The Kaplan-Meier plot shows survival of mice with treatment as indicated. Asterisks denote **p < 0.01, from Log- rank (Mantel-Cox) test. (E) C3H/HeNCrNarl mice were intravenously injected with 1 × 10^6^ cells of T315I-Bcr-Abl-32D cell. After 7 day of transplantation, mice were treated with vehicle (n = 7), or JMF4073 (5 mg/kg/time, n = 7) by intraperitoneal injection at 24 h interval for 14 days. The Kaplan-Meier plot shows survival of mice with treatment as indicated. The Kaplan-Meier plot shows survival of mice with treatment as indicated.

### ROS-induced replication stress and dTTP deficiency determine JMF4073 sensitivity

To understand the differential sensitivity to JMF4073, we then compared the levels of four dNTPs and replication stress in untransformed, WT- and T315I-Bcr-Abl-32D cells. The quantitation of four dNTPs revealed lower dNTP levels in WT-Bcr-Abl-32D cells than T315I-Bcr-Abl-32D and untransformed 32D cells, particularly the dTTP pool (Fig 3A). DNA fiber analyses were performed to evaluate replication stress. We found that the length of IdU- labeled replication track linking to CldU-labeling in WT-Bcr-Abl cells much shorter than those in T315I-Bcr-Abl- and untransformed 32D cells, indicating very prominent DNA replication stress present in WT-Bcr-Abl-32D cells but not the other two cell lines (Fig 3B). Treatment of WT-Bcr-Abl cells with N-acetyl cysteine (NAC), a ROS scavenger, increased the lengths of replication track, (Fig 3C). Importantly, NAC or GSH-MEE treatment caused WT-Bcr-Abl cells no longer vulnerable to JMF4073 treatment (Fig 3D), suggesting the determining role of ROS in replication stress and JMF4073 sensitivity. The cellular levels of GSH and dTTP were elevated by NAC treatment (Fig 3E-F). GSH and GSSG measurements showed that GSH level was significantly lower in WT-Bcr-Abl cells (Fig 3G). Since exogenous supplementation of dT and dC for 4 days also mitigated replication stress and DNA damages in WT-Bcr-Abl cells (Appendix Fig S2), these data suggest that nucleotide supply and redox status determine JMF4073 susceptibility.

**Figure 3.**
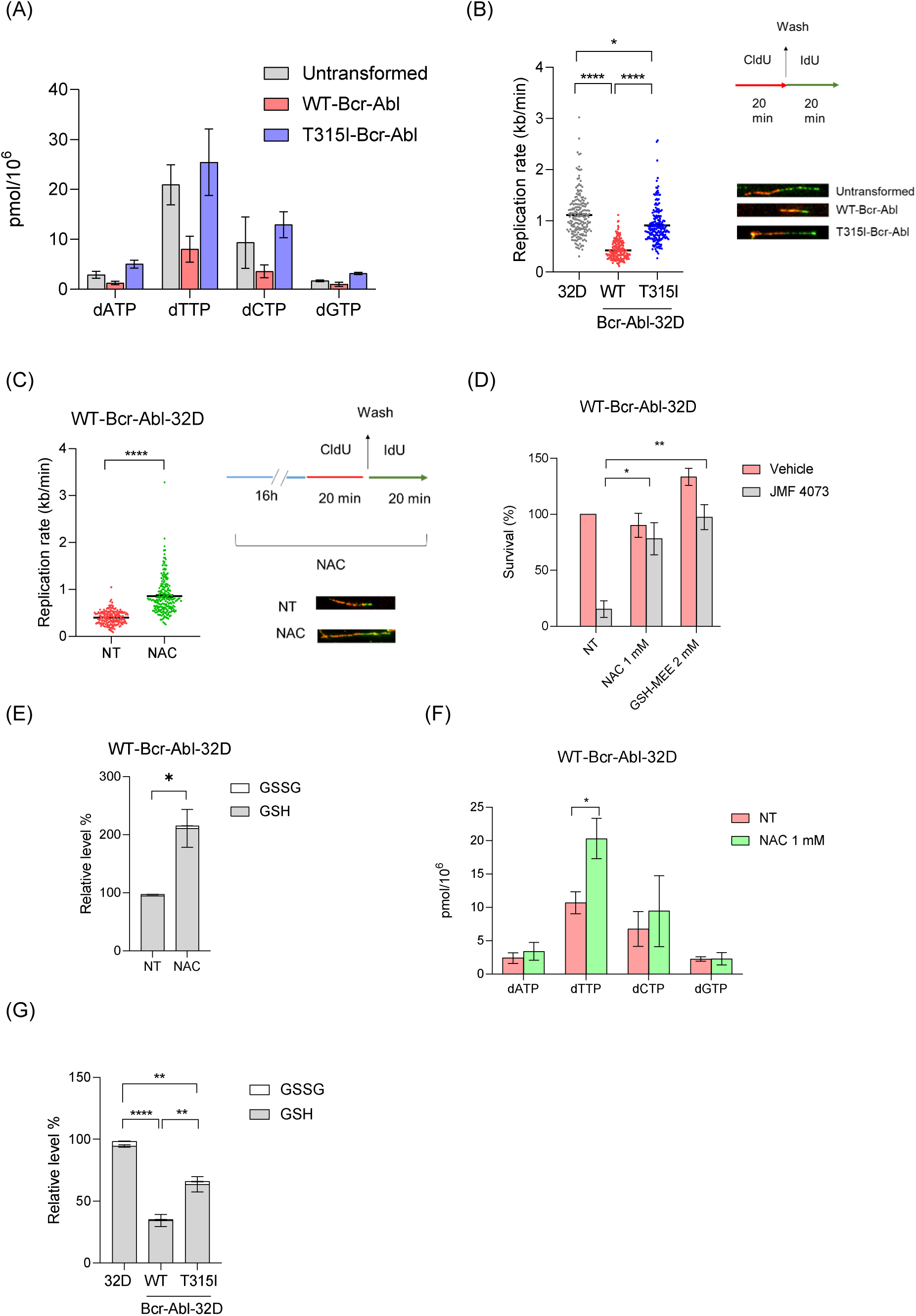
ROS-induced replication stress is involved in JMF4073 vulnerability. (A) Four dNTP levels in untransformed, WT-, T315I-Bcr-Abl-32D cells. Data are represented as means ± SEM, n > 3 biological replicates. (B) Replication rate in untransformed, WT-, and T315I-Bcr-Abl transformed 32D cells was measured by DNA fiber analysis (n = 200 fibers for each cell line) (*Left)*. DNA fiber labeling was performed by incubated with 5-chloro-2’-deoxyuridine (CldU, red) for 20 min followed by washout and incubation with 5-iodo-2’-deoxyuridine (IdU, green) (*Right*). The representative labeled fibers were shown below. Replication rate in kb/min was calculated by the measured length of IdU (green) linking to CldU (red) in μm with a conversion factor of 0.34 μm/kb and dividing by the duration of the labeling pulse. (C) Effect of ROS scavenger on replication stress. WT-Bcr-Abl-32D cells were treated with N-acetyl cysteine (NAC, 1 mM) for 16 h prior to DNA fiber analysis as indicated (*Right*). The DNA replication rate was determined by the length of IdU (green)-labeled fibers linking to CldU (red) (n = 200 fibers for each condition). (D) Redox-dependent JMF4073 sensitivity. WT-Bcr-Abl cells were treated with NAC (1mM) or GSH ethyl- ester (GSH-MEE, 2 mM) for 24 h, followed by the addition of JMF4073 (1 μM). After 3 days, viability was measured by CCK-8 assay. Data are represented as means ± SEM, n = 3 biological replicates. (E) WT-Bcr-Abl-32D cells treated with NAC (1 mM) for 24 h were harvested for GSH and GSSG determination. Data are represented as means ± SEM, n > 3 biological replicates. (F) The dNTP pools in WT-Bcr-Abl-32D cells after 1 mM NAC treatment for 24 h from 3 independent experiment. (G) GSH and GSSG levels in untransformed, WT-, T315I-Bcr-Abl-32D cells. Data are represented as means ± SEM, n > 3 biological replicates. Asterisks denote *p < 0.05, **p < 0.01, ***p < 0.001, or ****p<0.0001 from non-paired two-tailed Student’s t test.

### T315I-Bcr-Abl impacts mitochondrial regulation to change dNTP and ROS levels

We further asked why T315I-Bcr-Abl transformation did not cause prominent ROS-induced replication stress and dTTP deficiency as seen in WT-Bcr-Abl-32D cells. It should be mentioned the levels of enzymes, such as R1 and R2 subunits of RNR, TS, TMPK, and thymidine kinase 1(TK1) in nucleotide biosynthesis, in WT-Bcr-Abl were not less than those in T315I-Bcr-Abl cells (Appendix Fig S3). It has been shown that the activity of CAD, the key enzyme for de novo pyrimidine synthesis, is upregulated by S6K (Ben-Sahra, Howell et al., 2013, Robitaille, Christen et al., 2013), which is a downstream effector of Bcr-Abl signal. However, the major downstream signals of Bcr-Abl including Stat5, S6K and AKT, were similar in WT and T315I-Bcr-Abl-32D cells (Appendix Fig S4). Since RNR-mediated dNTP formation and TS-mediated dTTP formation require co-factor NADPH, we then compared the level of NADPH in these cells. The results showed lowest level of NADPH in WT-Bcr-Abl-32D (Fig 4A). Altogether, lower levels of GSH and NADPH suggest WT-Bcr-Abl cells under higher oxidative stress.

**Figure 4.**
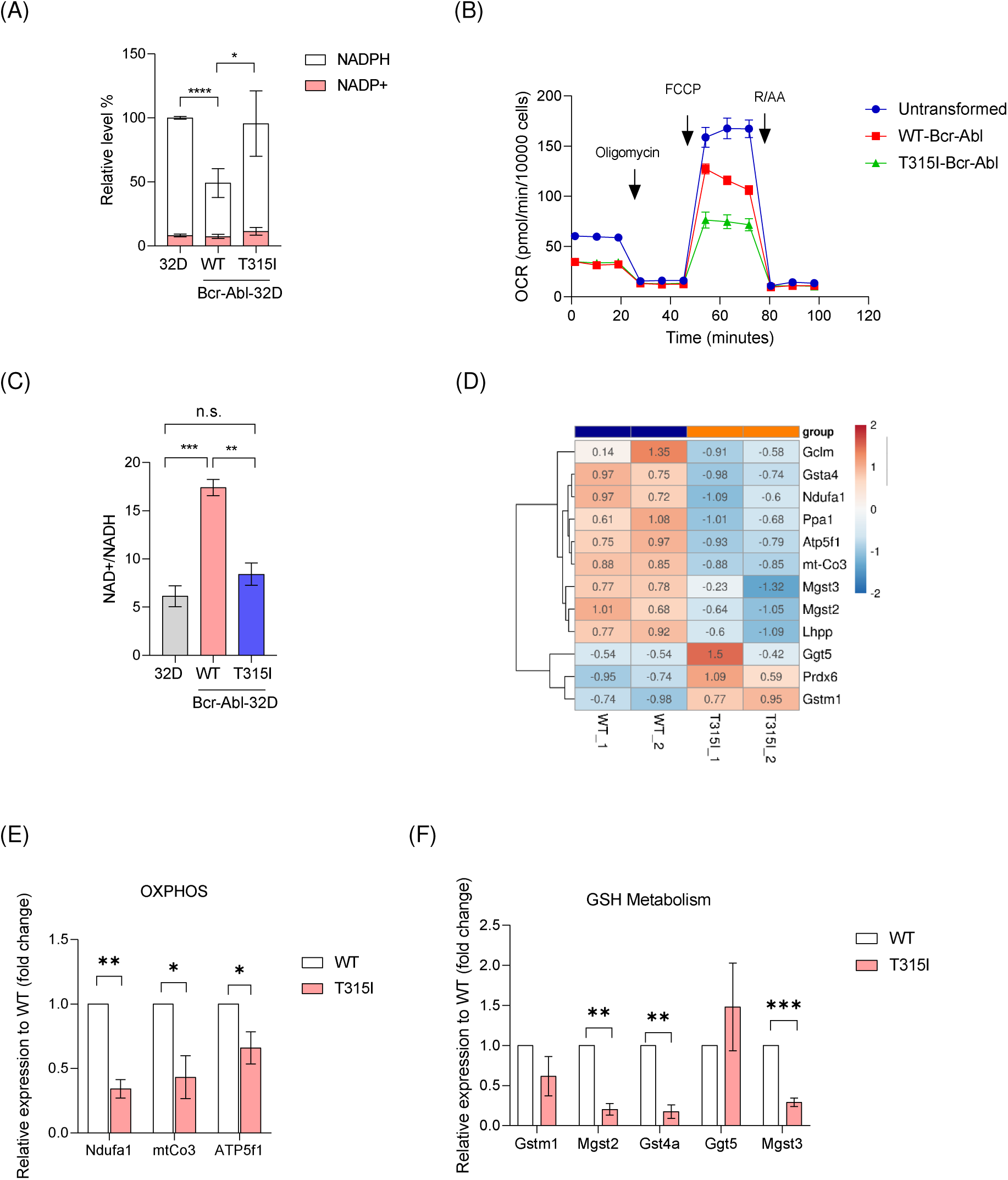
T315I mutation affects redox, energetic, and OXPHOS statuses. Untransformed, WT-, T315I-Bcr-Abl-32D cells as indicated were subjected to the analyses. (A) NADPH and NADP+ levels. Data are presented as relative to those in untransformed 32D cells from three independent experiments. (B) Oxygen consumption rate (OCR) measurement (n=3). (C) NAD+/NADH ratio was measured in untransformed, WT-, T315I-Bcr-Abl-32D cells. (D) Transcriptome analysis. RNA isolated from WT- and T315I-Bcr-Abl-32D cells were used for RNA-sequencing analysis. Heat map showing RNA-sequencing data for genes in OXPHOS and GSH metabolism, sorted by log_2_ Fold Change and adjust p-value <0.05. The values on heat map are built by using DESeq 2 on normalized gene read counts. (E-F) RT-qPCR of genes transcripts in OXPHOS and GSH metabolism identified in (D). The bar-plots show relative mRNA transcripts normalized to 18s rRNA primer sets. Data are represented as means ± SEM, n = 3 biological replicates. Asterisks denote *p < 0.05, **p < 0.01, or ***p< 0.001 from non-paired two-tailed Student’s t test.

Mitochondrion plays a central role in ROS production and the redox status. We then evaluated OXPHOS activity by oxygen consumption rate (OCR) analysis. As compared to untransformed 32D cells, the basal OCR and maximal respiration were significantly decreased in WT- and T315I-Bcr-Abl-transformed cells (Fig 4B), confirming Bcr-Abl-induced mitochondrial dysfunction. Of note, the level of maximal respiration was much lower in T315I- than that in WT-Bcr-Abl cells (Fig 4B). NAD+ is the major co-factors in the oxidation reactions of TCA cycle and fatty acid degradation in NADH production, which is subsequently oxidized to NAD+ by electron transfer chain reactions in mitochondria. NAD+/NADH ratio was higher in WT-Bcr-Abl cells (Fig. 4C), consistent with higher respiratory activity. To access whether T315I mutation affect the levels of genes that regulate redox and OXPHOS, we performed RNA sequencing analysis. Among the 1515 genes that were differentially expressed, 862 genes were upregulated, and 652 genes were downregulated in T315I-Bcr-Abl-32D cells (Appendix Fig S5A). The 10 most significantly enriched upregulated and down- regulated gene ontology (GO) terms were mainly related to migration and immunity (Appendix Fig S5B-S5C). Nevertheless, the expression of genes involved in the biological pathways related to anti-oxidative stress, ROS metabolism, and cofactors catabolic process were found significantly up-regulated in T315I-Bcr-Abl cells (Appendix Fig S5D). The analysis led us to find the differences in gene expression related to GSH metabolism and OXPHOS genes (Fig 4D; Appendix Fig S5E). RT-qPCR validated that the RNA transcript levels of Ndufa1, mt-Co3 and ATP5f1b, encoding the essential component of I, IV, and V, respectively, were significantly lower in T315I-Bcr-Abl cells (Fig 4E). The transcript levels of genes, Mgst2, Gsta4, and Mgst3, responsible for GSH conjugation were also lower in T315I-Bcr-Abl cells. These changes in RNA transcripts might contribute to reduced OXPHOS activity and higher GSH level in T315I-Bcr-Abl cells (Fig 4F).

We further measured mitochondrial DNA copy number, which showed higher level in WT-Bcr-Abl cells (Fig 5A). It has been reported that Sirt1, a NAD+ dependent deacetylase, is highly expressed in CML cells (Abraham et al., 2019, Li, Wang et al., 2012, Yuan, Wang et al., 2012). In addition, Sirt1 depletion delays the progression of CML (Abraham et al., 2019). It is well established that Sirt1 promotes the co- transactivation function of PGC1α, a mitochondria biosynthesis factor, by deacetylation (Lagouge, Argmann et al., 2006, Rodgers, Lerin et al., 2005). We then analyzed that the expression level of Sirt1, and found Sirt1 upregulation in WT- but not- T315I-Bcr-Abl-32D cells (Fig 5B). To know whether higher Sirt1 level contributes to the difference in mitochondrial DNA copy number and OCR, Sirt1 was knocked down in WT-Bcr-Abl-transformed cells by lentiviral infection of Sirt1 shRNA. Mitochondrial DNA copy number and maximal OCR were significantly decreased after Sirt1 knockdown in WT-Bcr-Abl cells (Fig 5C-5D), indicating Sirt1 upregulation responsible for higher mitochondrial copy number and maximal OXPHOS.

**Figure 5.**
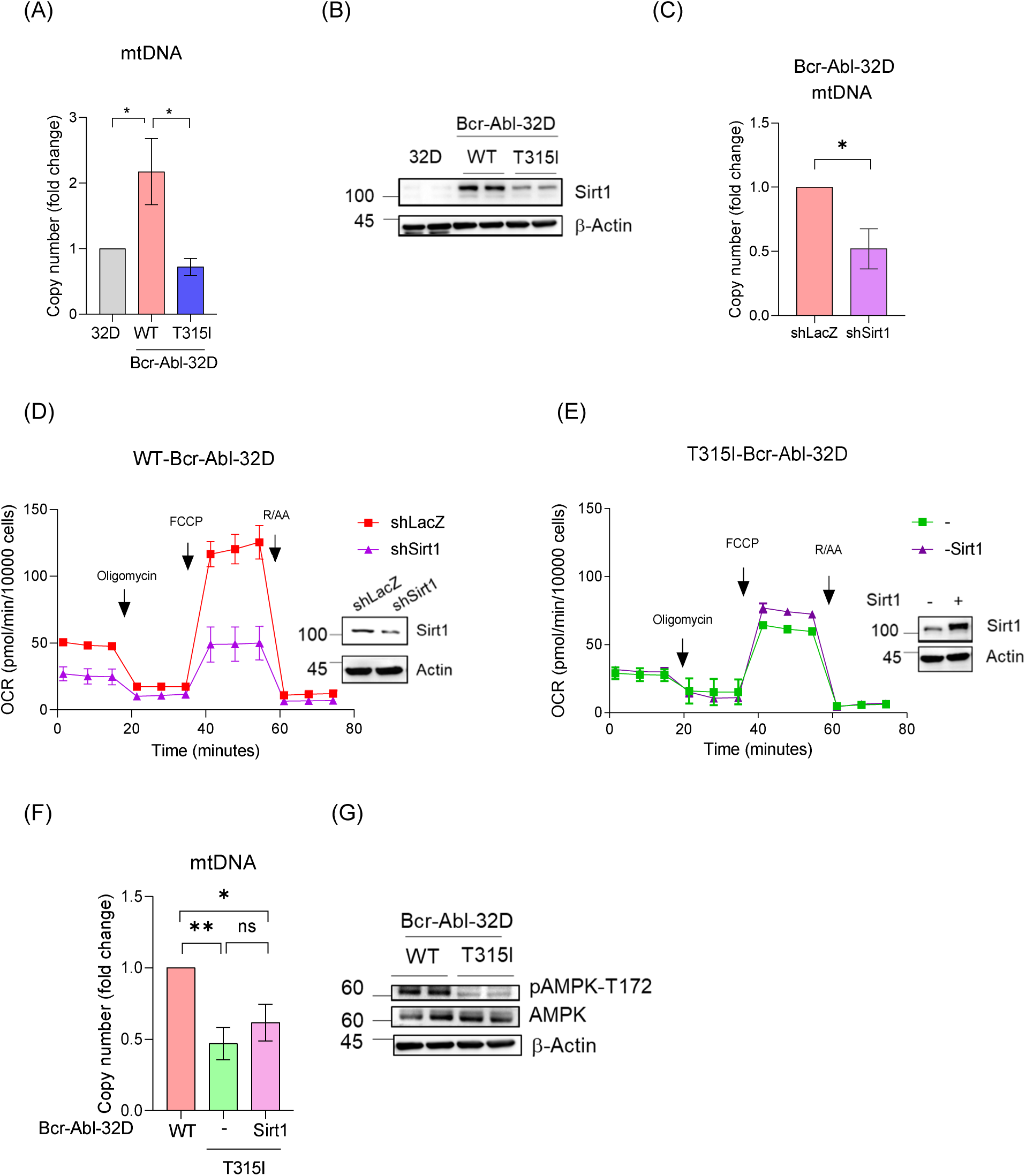
Sirt1 regulates OXPHOS in WT- but not T315I-Bcr-Abl cells. (A) The mitochondria copy number. DNA were isolated for qPCR analysis using mt- 16s rRNA primer set and normalized by nuclear DNA using mouse β-globin primer set. Data are represented as means ± SEM from three independent experiments. (B) The expression level of Sirt1 in untransformed, WT-, T315I-Bcr-Abl-32D cells. Western blots of two independent sets of samples. (C-D) WT-Bcr-Abl-32D cells were infected with shLacZ and shSirt1 lentivirus for (C) mtDNA copy number and (D) OCR measurement analysis. Data are represented as means ± SEM from 3 independent experiments. (E-F) T315I-Bcr-Abl-32D cells were infected with control or Sirt1retrovirus for (E) OCR measurement, and (F) measuring mtDNA copy number. Data are represented as means ± SEM from 3 independent experiments. Asterisks denote *p < 0.05, or **p < 0.01 from non-paired two-tailed Student’s t test. (G) The comparison of AMPK phosphorylation in WT- and 315I-Bcr-Abl-32D cells by Western blot.

However, in T315I-Bcr-Abl-32D cells, overexpression of Sirt1 was unable to increase mtDNA copy number nor OXPHOS activity (Fig 5E-F). It has been suggested that AMPK activates Sirt1 in metabolic regulation (Lan, Cacicedo et al., 2008, Ruderman, Xu et al., 2010). We found the level of active AMPK obviously very low in T315I-Bcr-Abl cells (Fig 5G). It is known that ATP competes AMP in AMPK binding to prevent activation (Herzig & Shaw, 2018, Xiao, Sanders et al., 2013, Xiao, Sanders et al., 2011). Accordingly, high ATP level found in T315I-Bcr-Abl cells might contribute to the deficiency in AMPK activation (Fig 2C). The cellular context of high ATP and less active AMPK suggests higher energy state in T315I cells, which might disable Sirt1 in the stimulation of mitochondrial biogenesis. Overall, these data show that T315I mutation alters Bcr-Abl-regulated mitochondrial function.

### Reductive metabolism of glutamine in TCA cycle and GSH synthesis in T315I-Bcr-Abl cells

To detail the metabolic changes, metabolomic analysis by mass spectrometry in WT- and T315I-Bcr-Abl cells were performed. The results revealed higher levels of amino acids, nucleoside metabolites, and glycolytic intermediates in T315I-Bcr-Abl cells, confirming the metabolic changes caused by T315I mutation (Appendix Fig S6). It is well established that in cancer cells, glutamine replenishes TCA cycle intermediates via reductive carboxylation or oxidative carboxylation to feed biosynthetic pathways (Altman, Stine et al., 2016, Hatzivassiliou, Zhao et al., 2005, Kovacevic, 1971, Smith, Schafer et al., 2016, Vander Heiden, Cantley et al., 2009) and provides GSH synthesis (Fig 6A). WT- and T315I-Bcr-Abl-32D cells incubated with (U)-^13^C-glutamine were analyzed to trace the metabolic fates of glutamine. The analyses showed significant increases in both m+5-citrate and m+3-malate in T315I-Bcr-Abl cells as compared to those in WT-Bcr-Abl cells, while no significant difference in both m+4-citrate and m+4- malate (Fig 6B). In addition, an obvious increase in m+5 GSH was observed in T315I- Bcr-Abl cells (Fig 6B). Altogether, these data indicate that T315I-Bcr-Abl cells have more glutamine flux to GSH synthesis and reductive carboxylation in the TCA cycle. Thus, a metabolic network programmed by Bcr-Abl transformation is shifted towards the reductive status by T315I mutation.

**Figure 6.**
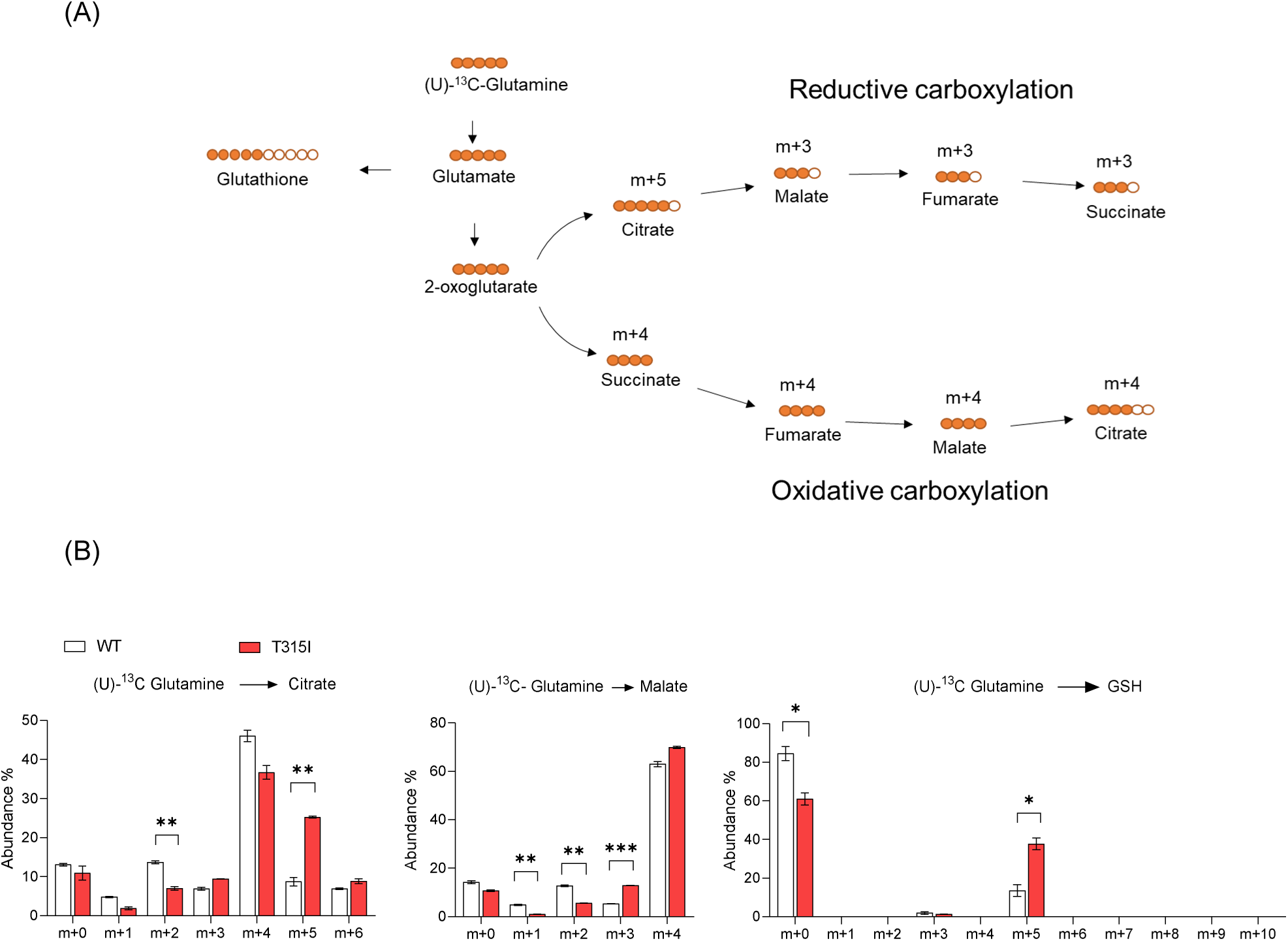
T315I mutation causes reductive carboxylation. (A) Schematic illustration of the metabolic path U-^13^C glutamine incorporation into the TCA cycle and GSH. Red indicates ^13^C label, and ^12^C non-label by white. (B) Traced isotopologue abundance in WT- and T315I-Bcr-Abl-32D cells after incubated with 4 mM U-^13^C glutamine medium for 2 h. Data are presented as means ± SEM, n > 3 biological replicates. Asterisks denote *p < 0.05, or **p < 0.01, from non-paired two-tailed Student’s t test.

### Converting T315I-Bcr-Abl-32D cells vulnerable to JMF4073 by blocking MPC

Next, we wish to switch T315I-Bcr-Abl cell more oxidative to increase JMF4073 sensitivity. Three metabolic targets including glucose-6-phosphate dehydrogenase, glutaminase, and mitochondrial pyruvate carrier that regulate the reductive status were chosen. Treatment of T315I-Bcr-Abl cells with 6-AN, an inhibitor of glucose-6- phosphate dehydrogenase, did not increase JMF4073 sensitivity (Appendix Fig S7A- S7B). Inhibition of glutaminase by BPTES in combination with JMF4073 suppressed the growth of T315I-Bcr-Abl- and 32D myeloid progenitor cells similarly, indicating poor selectivity (Appendix Fig S7C). It has been shown that blocking mitochondrial pyruvate carrier (MPC) by UK-5099 diverts glutamate metabolism away from GSH biosynthesis (Tompkins, Sheldon et al., 2019). Remarkably, UK-5099 treatment significantly sensitized T315I-Bcr-Abl but not 32D cells to JMF4073 (Fig 7A). Importantly, treatment of T315I-Bcr-Abl cells with UK-5099 decreased GSH level (Fig 7B) with a coincident reduces in NADH level and increases in NAD+/NADH ratio (Fig 7C-7D), indicating increasing oxidative status in T315I-Bcr-Abl transformed cells. In agreement, replication stress was induced by UK-5099 treatment (Fig 7E). Of note, the combination of UK-5099 and JMF4073 depleted dTTP and dCTP pools (Fig 7F). Finally, in vivo therapy of UK-5099 with JMF4073 was performed. We transplanted 32D Bcr-Abl-T315I/EGFP cells to C3H/HeNCrNarl mice. After 7 days, these mice were treated with a daily intraperitoneal injection of UK-5099 alone or UK-5099 in combination with JMF4073 for 14 days (Fig 7G-7H). The survival of T315I leukemia mice was not affected by UK-5099 therapy (Fig 7G). As a contrast, the combination of UK-5099 and JMF4073 therapy prolonged the survival (Fig 7H). Moreover, mice blood analysis revealed marked decreases in the population of Bcr-Abl-T315I/EGFP + cells by the combination of JMF4073 with UK-5099 treatment (Fig 7I), so as the spleen size (Fig 7J).

**Figure 7.**
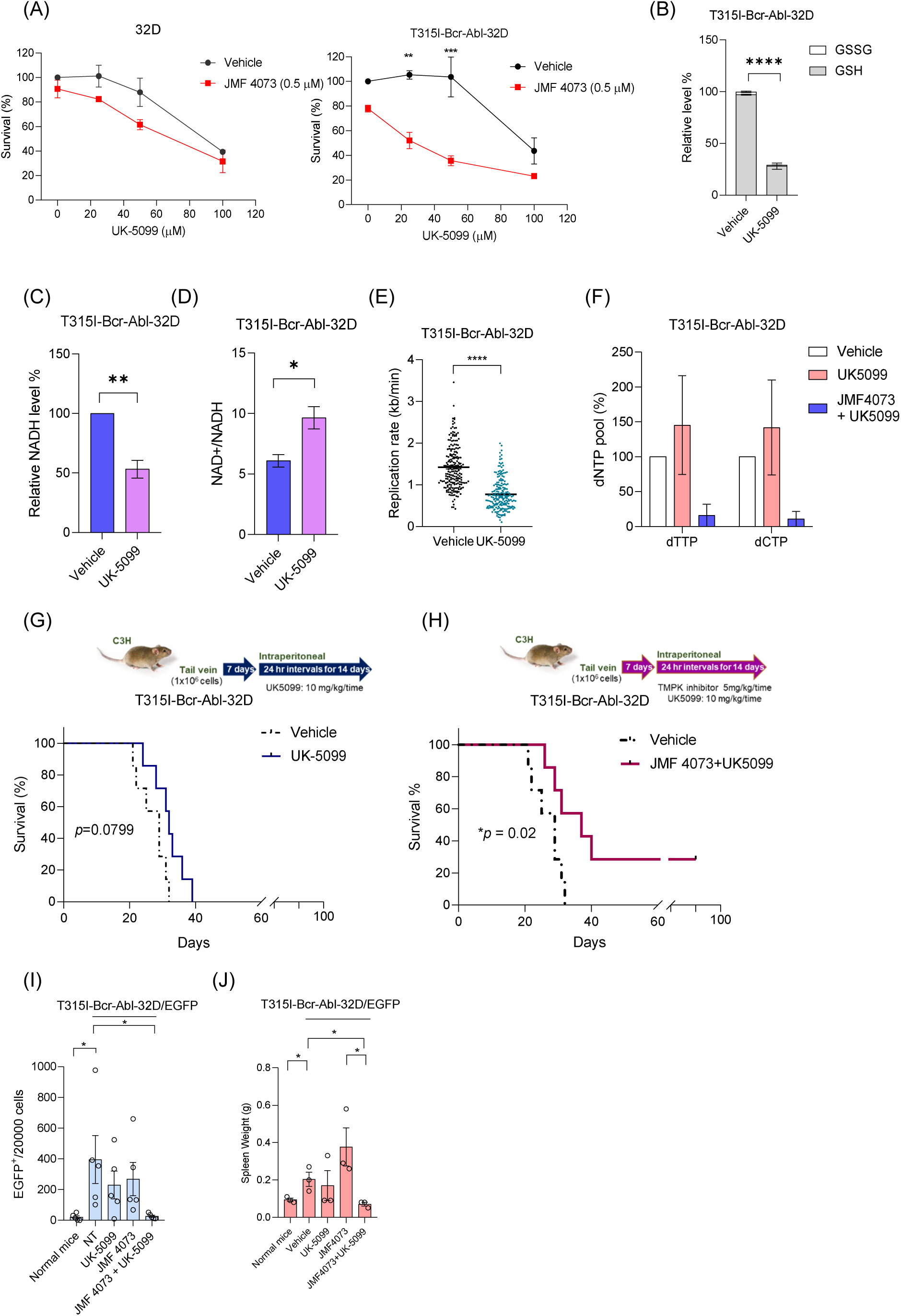
The synthetic effect of JMF4073 and UK-5099 in eradicating T315I Bcr- Abl-32D cell. (A) Untransformed or T315I-Bcr-Abl-32D cells were pre-treated with UK-5099 for 24 h, followed by the addition of vehicle or JFM4073. The viability of these cells was assayed after incubation for 3 days. (B-C) T315I-Bcr-Abl-32D cells were treated with UK-5099 (50 μM) for 24h. These cells were harvested for the analyses of (B) GSH and GSSG levels, (C) NADH level and (D) NAD+/NADH ratio. Data are presented as means ± SEM, n > 3 biological replicates. Asterisks denote *p < 0.05, or ****p<0.0001 from non-paired two-tailed Student’s t test. (E) The effect of UK-5099 on replication rates. T315I-Bcr-Abl-32D cells were pre-treated with UK- 5099 (50 μM) for 24h, followed by DNA fiber analysis as indicated. Data were from the length of labeled DNA fibers n = 200. Asterisks denote ****p<0.0001 from non- paired two-tailed Student’s t test. (F) The relative levels of dTTP and dCTP in response to UK-5099, and the combination of UK-5099 and JMF4073. T315I-Bcr-Abl-32D cells were pre-treated with or without UK-5099 (50 μM) for 16 h prior to the incubation with vehicle or JMF4073 (10 μM) for another 6 h. Data are presented as means ± SEM, n > 3 biological replicates. Asterisks denote *p < 0.05 from non-paired two-tailed Student’s t test. (G-H) The workflow of mouse leukemia induction and in vivo drug treatment: C3H/HeNCrNarl mice were intravenously injected with 1 × 10^6^ cells of T315I-Bcr- Abl-32D/EGFP. After 7 days of transplantation, mice were treated with vehicle (n = 7), (G)UK-5099 (10 mg/kg/time, n = 7) or (H) JMF4073 (5 mg/kg/time) + UK-5099 (10 mg/kg/time) (n=7) by intraperitoneal injection at 24 h interval for 14 days. The Kaplan-Meier plot shows survival of mice with treatment as indicated. Asterisks denote *p < 0.05, from Log-rank (Mantel-Cox) test. (I) Flow cytometry analysis of the leukemic progression in vehicle, UK-5099, JMF4073, or JMF4073 + UK-5099 treated CML mice. The number of EGFP+ cells in mice with T315I-Bcr-Abl-induced CML was determined on day 30 after transplantation (n=3). (J) Spleen weights of T315I-Bcr-Abl-32D/EGFP+ bearing mice treated with vehicle, UK-5099, JMF4073, or JMF4073 combined with UK-5099 (n=3). Data are represented as means ± SEM, n > 3 biological replicates. Asterisks denote *p < 0.05 from non-paired two-tailed Student’s t test.

## Discussion

In this study, we showed ROS-induced replication stress and dNTPs pool deficiency in WT-Bcr-Abl-transformed myeloid cells, making these cells vulnerable to pyrimidylate kinases inhibition by JMF4073. In a murine CML model bearing WT-Bcr-Abl-32D cells, JMF4073 treatment prolonged mice survival. Contrarily, T315I-Bcr-Abl- transformation did not cause prominent ROS-induced replication stress nor dNTPs deficiency in myeloid cells. Therefore, T315I-Bcr-Abl-32D cells were less sensitive to JMF4073, and the survival of CML mice bearing T315I mutation was not significantly affected by JMF4073 therapy. However, blocking mitochondrial pyruvate carrier by UK-5099 was able to decrease GSH level and to induce replication stress in T315I-Bcr-Abl cells, thereby converting these cells sensitive to JMF4073. Consistently, the survival of CML mice bearing T315I mutation was prolonged by the combinatory therapy of UK-5099 and JMF4073 for two weeks. The mechanism for this metabolic targeting is summarized as depicted (Appendix Fig S8). Since myeloid progenitor cells were not so sensitive to JMF4073, UK-5099, or the combined treatment by JMF4073 with UK-5099, our data suggest the potential of combining JMF4073 and UK-5099 for eradicating T315I-Bcr-Abl CML with little toxicity to the normal myeloid progenitor cells.

Consistent with the previous report, Bcr-Abl-mediated transformation causes a marked reduction of mitochondrial respiration capacity in 32D progenitor cells. Intriguingly, as compared to T315I-Bcr-Abl, WT-Bcr-Abl cells had higher OXPHOS activity and NAD+/NADH ratio with increased mitochondrial DNA copy number. These data indicate higher oxidative load in WT-Bcr-Abl cells. Previous findings have demonstrated that Sirt1 is upregulated in CML cancer stem cell and is responsible for stimulating OXPHOS function (Abraham et al., 2019, Huang, Lu et al., 2021, Tang, 2016). This study uncovered that the level of Sirt1 was clearly upregulated in WT- but not T315I-Bcr-Abl cells. Sirt1 knockdown in WT-Bcr-Abl cells did reduce OXPHOS activity and mtDNA copy number. However, enforced expression of Sirt1 in T315I- Bcr-Abl-32D cells was insufficient to boost OXPHOS activity nor the mtDNA copy number. Since the expression of a number of genes in OXPHOS reactions was decreased in T315I-Bcr-Abl cells, Sirt1 overexpression is probably insufficient to upregulate OXPHOS function. In the meanwhile, T315I-Bcr-Abl cells had higher level of ATP and little AMPK activation, thus restraining PGC-1α activation. Sirt1 upregulation in CML has been shown to cause p53 inactivation by de-acetylation (Li et al., 2012). Theoretically, the cooperation of p53 suppression with replication stress drives genome evolution to acquire mutations that are selected by TKI. Our findings suggest that T315I mutation in Bcr-Abl not only confers TKI resistance, but also rewires Bcr-Abl-regulated metabolism that collaterally reduces ROS-induced replication stress.

In this study, we found the association of higher reductive status in T315I cells with a significant increase in reductive carboxylation of α-ketoglutarate from glutamate in the TCA cycle. Moreover, the glutamine isotope flux analysis further showed the increase of GSH synthesis in T315I cells. These metabolic changes evoke the question of what causes these alterations. Of note, despite of T315I mutation in the ATP binding pocket of Bcr-Abl kinase domain, the overall tyrosine phosphorylation pattern and multiple proliferation signals, such as PI3K/AKT/mTOR and STAT5 pathway, were similarly stimulated in WT- and T315I-Bcr-Abl-transformed 32D cells. It is possible that T315I and WT Bcr-Abl have subtle differences in substrates, which relay pleiotropic signals that not only control metabolic regulators but also gene transcription as indicated by different gene profiles by RNA sequencing analysis. Relevant to the findings of this study, the transcript levels of three members in ETC complex I, IV and V and several glutathione transferases were lower in T315I cells, which might partly explain the reduced OXPHOS function and higher level of GSH. Nevertheless, it still remains to investigate what substrates are differentially tyrosine phosphorylated by WT- and T315I-Bcr-Abl that directly determine the changes of the energetic metabolism and redox.

Importantly, this study showed that blocking mitochondrial pyruvate carrier by UK-5099 effectively increased oxidative load in mitochondria as indicated by reducing GSH and increasing NAD+/NADH ratio in T315I-Bcr-Abl cells. The combination with JMF4073, UK-5099 treatment indeed depleted dTTP and dCTP pools, therefore suppressing the in vitro and in vivo growth of T315I cells. In conclusion, this study uncovered the reductive shift in metabolism by T315I mutation in Bcr-Abl transformed myeloid cells, and revealed a new compound, JMF4073, which inhibits TMPK and CMPK with higher potency and solubility than the previously reported TMPK inhibitor. We proposed that JMF4073 treatment in WT-Bcr-Abl cells and UK-5099-treated T315I-Bcr-Abl cells brought down dTTP and dCTP pools below the threshold required for overcoming replication stress and repairing DNA damages for growth, thereby eliminating CML cells.

## Material and methods

### Materials

HPLC-grade methanol, ethanol, Triton-X-100, 2-propanol, imatinib, UK-5099, adenosine triphosphate (ATP), thymidine monophosphate (TMP), cytidine monophosphate (CMP), lactate dehydrogenase, pyruvate kinase, NADH, phosphoenolpyruvate, U-^13^C-glucose, and U-^13^C-glutamine were from Sigma Aldrich. Ni-NTA resin was purchased from Qiagen. Glutathione sepharose 4B and thrombin were from GE (Cytiva). Antibodies: anti-TMPK and anti-CMPK were prepared as described previously (Ke, Kuo et al., 2005); anti-Abl, anti-Stat5, anti-phospho-Stat5 (Tyr694), anti-Akt, anti-phospho-Akt (Ser473), anti-phospho-Akt (Thr308), anti-AMPK, anti-phospho-AMPK (Thr172), anti-Erk1/2, anti-phospho-Erk1/2 (Thr202/Tyr204), and anti-Sirt1 were obtained from Cell Signaling; anti-β-actin, anti-mouse IgG (HRP), anti-rabbit IgG (HRP), and anti-goat IgG (HRP) were from GeneTex; anti-RRM1, anti-RRM2, and anti-phospho-Tyr99 were from Santa Cruz.

### Cell lines, cell culture and virus infection

Murine WEHI, 32D (32Dcl3), WT and T315I-BcrAbl-transformed 32D cells were kindly provided by Dr. RB Arlinghaus (The University of Texas MD Anderson Cancer Center, Houston, TX, USA.) and maintained in RPMI 1640 medium supplemented with 10% heat-inactivated fetal bovine serum, 1% Streptomycin/Penicillin,1mM sodium pyruvate, and 10 mM HEPES. The growth medium for 32D cells additionally contained 10% conditioned medium from the WEHI-3B cell culture, as a source of murine interleukin-3 (IL3). Lentivirus and retrovirus infection are described in supplementary methods.

### Chemistry

Compound JMF4073, *Tert*-butyl 4-(2-(5-fluoro-3-oxobenzo[*d*]isothiazol-2(3*H*)-yl) acetyl) piperazine-1-carboxylate, was synthesized using 2,5-difluorobenzonitrile as the starting materials. See supplementary methods for synthetic schemes and procedures with NMR spectrums.

### Purification of recombinant proteins

pGEX-2T-hTMPK and pET28c-CMPK were each transformed into an *E. coli* BL21 strain. A single clone was inoculated in 5 ml of LB broth and cultured overnight at 37°C. 1 mL of the overnight culture was diluted to 100 mL, and cell growth was continued at 37 °C until the OD_600_ reached 0.5. For hTMPK and hCMPK induction, isopropyl β-d-1-thiogalactopyranoside (IPTG) was added to a final concentration of 0.5 mM, and the culture was incubated for another 3.5 h at 37 °C. For hTMPK purification, bacteria were harvested by centrifugation and re-suspended in 4 mL of lysis buffer containing 1% Triton X-100, 50 mM Tris-HCl, pH 7.5, 150 mM NaCl, 1 mM DTT, 1 mM PMSF, and 1 mM protease mixture. After sonication and centrifugation, 400 μL of GSH-Sepharose beads was added to the clarified cell lysate and incubated with gentle shaking at 4 °C for 2 h. Beads were washed five times with a buffer containing 0.5% Triton X- 100, 50 mM Tris-HCl, pH 7.5, 150 mM NaCl, 5 mM EDTA, 1 mM DTT, 1 mM PMSF, and 1 mM protease mixture. The GST-tag was then cleaved from the target proteins by thrombin.

For hCMPK purification, bacterial lysates were prepared in 4 ml of lysis buffer containing 1% Triton X-100, 50 mM Tris-HCl, pH 7.5, 150 mM NaCl, 1 mM TCEP, 1 mM PMSF, and 1 mM protease mixtures. After sonication and centrifugation, 200 μL of Ni-NTA resin was added to the clarified lysate and incubated with gentle shaking at 4 °C for 2 h. Beads were washed five times with a buffer containing 0.5% Triton X- 100, 50 mM Tris-HCl, pH 7.5, 150 mM NaCl, 5 mM imidazole, 1 mM TCEP, 1 mM PMSF, and 1 mM protease mixture. The His-CMPK protein was then eluted from Ni-NTA resin by elution buffer containing 50 mM Tris-HCl, pH 7.5, 150 mM NaCl, and 500 mM imidazole.

### Oxygen consumption measurement

XFe24 cell culture microplates were coated with poly-L-lysine (Sigma) or Cell-Tak (Corning) prior to seeding cells for the oxygen consumption rate (OCR) measurement. Cells (4×10^4^) were washed twice with PBS and re-suspended with 175 μL of assay medium containing bicarbonate-free RPMI1640 (Sigma-Aldrich) supplemented with 25 mM glucose, 1 mM sodium pyruvate, 2 mM glutamine, and 2% FBS cells, for seeding in XFe24 plate. After cell attachment, 500 μL of assay medium was added to each well, followed by incubation for 30 minutes at 37°C in a non-CO_2_ incubator. OCR was assayed in a Seahorse XF-24 extracellular flux analyzer by the sequential addition of 1 μM oligomycin (port A), 1 μM carbonyl cyanide-p-trifluoromethoxyphenylhydrazone (FCCP, port B), 1 μM rotenone and 1 μM antimycin A (port C).

### NADH-coupled enzymatic assay

Activity assay was measured using the spectrophotometric method by coupling ADP formation to NADH oxidation catalyzed by pyruvate kinase and lactate dehydrogenase. Each reaction contained 0.4 μg TMPK or 0.3 μg CMPK in the mixture containing 100 mM Tris-HCl, pH 7.5, 100 mM KCl and 10 mM MgCl_2_, 500 μM phosphoenolpyruvate, 250 μM NADH, 4 unit of pyruvate kinase and 5 unit of lactate dehydrogenase together with 500 μM ATP, 500 μM dTMP or dCMP. Reactions were incubated at 25 ℃ followed by the change in NADH measurement at 340 nm absorption.

### Cell viability assay

Cells were seeded at the density of 1×10^3^ cells/well in 96-well microplate. After 3 days, cell viability was determined using the CCK-8 cell viability. In brief, 20 μL of CCK-8 reagent were added to each well and incubated at 37℃ for at least 30 minutes. The viability was measured by using Tecan Spark 10 microplate reader at 450 nm according to the manufacturer’s instructions.

### Viral infection

For lentiviral production, HEK293T cells were co-transfected with 4 μg Δ8.91, 1 μg pVSVG, 5 μg of lentivector by using 30 μL of Turbofect (Thermo) in 1 mL of Opti-MEM according to the manufacturer’s instructions. HEK293T-PlatA cells were transfected with 10 μg of retrovector using 30 μL of Turbofect in 1 mL of Opti-MEM for retroviral production. Viral supernatants were collected 48 h later and used to infect WT-or T315I-Bcr-Abl-32D cells in the presence of 8 μg/ml polybrene. Cells were then selected and maintained in culture media containing 1 μg/mL of puromycin.

### NADH and NADPH measurement

NAD+/NADH and NADPH/NADP+ measurements were performed by enzymatic cycling reaction assay kits (NADH-Glo and NADPH-Glo, Promega). 4×10^4^ cells were resuspend in 50 μL of PBS and were lyzed by adding 50 μL of NaOH-DTAB buffer containing 1% w/v dodecyltrimethylammonium bromide (DTAB) and 0.2N NaOH. The lysates were split into 2 samples: one treated with acid solution for NAD+ or NADP+ measurement; the other treated with alkaline solution to detect NADH or NADPH. For acid-treated sample, 25 μL of 0.4 N HCl was added into 50 μL NaOH-DTAB treated cells, and incubated at 60°C for 15 minutes. After incubation, 25 μL of 0.5 M Trizma base were added to the acid-treated samples for neutralization. For alkaline-treated samples, 50 μL of HCl/Tris solution (0.4 N HCl: 0.5 M Trizma base 1:1 v/v) were added to the 50 μL of NaOH-DTAB samples. 25 μL of each acid-treated and alkaline-treated samples were transferred to a clean white 96-well plate. After adding 25 μL of NADH-Glo mixtures (Promega) or NADPH-Glo mixture (Promega), the plate was incubated at room temperature for 30 minutes and luminescence was obtained determined by a Tecan Spark 10 microplate reader.

### GSH/GSSG measurement

GSH/GSSG measurement was performed by a glutathione S-transferase (GST) enzyme coupled reaction using (GSH/GSSG Glo kit, Promega). Cells (2×10^4^) suspended in 25 μL of PBS were incubated with 25 μL of Glutathione lysing reagent. For GSSG measurement, N-Ethylmaleimide (NEM) was added into lysing reagent and incubated for 5 minutes to generate NEM-Glutathione, which is not the substrate of GST. NEM-treated and non-treated samples were incubated with 50 μL of GST reaction mixtures for 30 min at room temperature. 25 μL of each were then transferred to a fresh white 96-well plate containing 25 μL of GSH-Glo mixtures (Promega). Luminescence was determined using a Tecan Spark 10 microplate reader

### Metabolomics analysis

1 ×10^6^ cells were seeded in a 10-cm dish for steady-state metabolomics analysis. After 48 h, cells were washed twice with cold PBS and were than extracted with 80 % methanol at 2×10^6^ cells/mL. 4-Chloro-DL-phenylalanine (Fenclonine, Sigma) was added as internal control at the final concentration of 50 μg/mL methanol extracts.

Cells in 10-cm dishes were washed twice for metabolite tracing by glucose-free or glutamine-free RPMI-1640 medium. For glucose tracing, 25 mM of U-^13^C-glucose was added into glucose-free RPMI-1640 medium together with 10% v/v HI-FBS, 1 mM HEPES and 1 mM sodium pyruvate for 1 h. For glutamine tracing, 4 mM of U-^13^C- glutamine was added to the glutamine-free RPMI-1640 medium with 10% v/v HI-FBS, 1 mM HEPES, and 1 mM sodium pyruvate . After 2h incubation, cells were washed twice with cold PBS and were extracted with 80% ice-cold methanol at 2×10^6^ cells/mL. Samples were incubated at -80℃ for overnight. After centrifugation, supernatants were transferred to fresh tubes and evaporated using a speed-vac.

Agilent 1290 Infinity II ultra-performance liquid chromatography (UPLC) system (Agilent Technologies, Palo Alto, CA, USA) coupled online to the Dual AJS electrospray ionization (ESI) source of an Agilent 6545 quadrupole time-of-flight (Q-TOF) mass spectrometer (Agilent Technologies, Palo Alto, CA, USA) was used for the analysis. The ACQUITY UPLC BEH amide column (1.7 μm, 2.1 × 100 mm, Waters Corp., Milford, MA, USA) at 40℃ was employed for sample separation. The mobile phase was composed of double-distilled water (eluent A) and 10% double-distilled water in acetonitrile (eluent B) eluted with 15 mM ammonium acetate and 0.3% NH_3_·H_2_O. The flow rate was 300 μL/min, and the sample injection volume was 2 μL. The instrument was operated in positive and negative full-scan mode, collected from a m/z of 60–1500. The MS operating conditions were optimized as follows: Vcap voltage, 3.5 kV; nozzle voltage, 1 kV for negative mode and 0 V for positive mode; nebulizer, 45 psi; gas temperature, 200 ℃; sheath gas temperature, 300 ℃; sheath gas flow (nitrogen), 8 L/min; drying gas (nitrogen), 10 L/min. The chromatogram acquisition, mass spectral peaks detection, and waveform processing were performed using Agilent Qualitative Analysis 10.0 and Agilent Profinder 10.0 software (Agilent, USA).

### Mouse CML induction and in vivo therapy

The animal studies were approved by the biosafety committee at National Taiwan University and conformed to the national guidelines and regulations. Female C3H/HeNCrNarl mice were used at 6-8 weeks of age (National Laboratory Animal Center, Taiwan). WT-Bcr-Abl-32D-EGFP+ cells (5 × 10^5^) suspended in 200 μL of HBSS were injected into each mouse through the tail vein. After 48h of transplantation, mice were treated with vehicle (DMSO), or JMF4073 (5 mg/kg body weight) by intraperitoneal injection at 24 h interval for 14 days. Injected mice were monitored for peripheral blood (PB) and counted by Taiwan Mouse Clinic-National Phenotyping Center, National Research Program for Genomic Medicine (NSC). For T315I-Bcr-Abl CML mice, T315I-Bcr-Abl-32D-EGFP+ cells (1×10^6^) suspended in 200 μL of HBSS were tail-vein-injected. After 7 days of transplantation, mice were treated with vehicle (DMSO), UK-5099 (10 mg/kg body weight), JMF4073 (5 mg/kg body weight), or UK- 5099 combined with JMF4073 by intraperitoneal injection at 24 h interval for 14 days. For monitoring CML progress, mice blood was collected in anticoagulation tubes by submandibular blood collection, followed by lysing red blood cells with RBC lysis buffer (Invitrogen). After washing, the remaining cells were suspended in 500 μL of HBSS and subjected to flow cytometry analysis (FACScalibur, BD Bioscience) with CellQuest software. The number of EGFP+ cells were collected from 20,000 single-cell events.

### Measurement of dNTP pools by RCA-qPCR assay

Cells (10^6^) were extracted with 1 ml of ice-cold 80% methanol at -80 ℃ for 16 hr, followed by heating at 95℃ for 3 minutes. The methanol extracts were centrifuged (16,000 x g, 30 min) to remove cell debris. For chloroform extraction, same volume of chloroform was added to the methanol extract and vortexed. After phase separation, upper phase was collected and vacuum-dried. The dried residues were dissolved in water, in which 1×10^4^ cells/ 5μL were used for assay according to a new method described previously (Wang, Hsin et al., 2021). Briefly, RCA reactions were performed in a mixture (10 μL) containing 50 mM Tris-HCl, 10 mM MgCl_2_, 10 mM (NH_4_)_2_SO_4_, 4 mM DTT, 0.1% Tween-20, pH 7.4, 2 units of phi29 DNA polymerase (New England Biolabs), three not to quantified deoxynucleoside triphosphate mix (20 μM), and 1 attomol of the annealed template-primer (the sequence listed below). The reaction was incubated at 37°C for 1 h and terminated by heating at 65°C for 10 min. The RCA products were determined by the subsequent addition of another 15 μL of qPCR reaction mixture (PCR Biosystem).

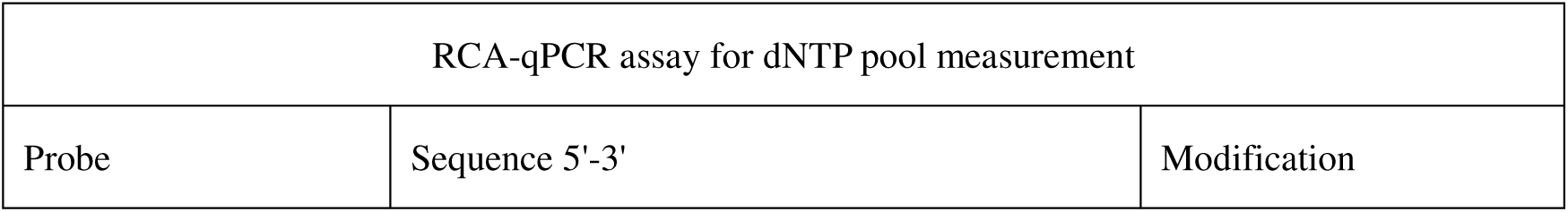
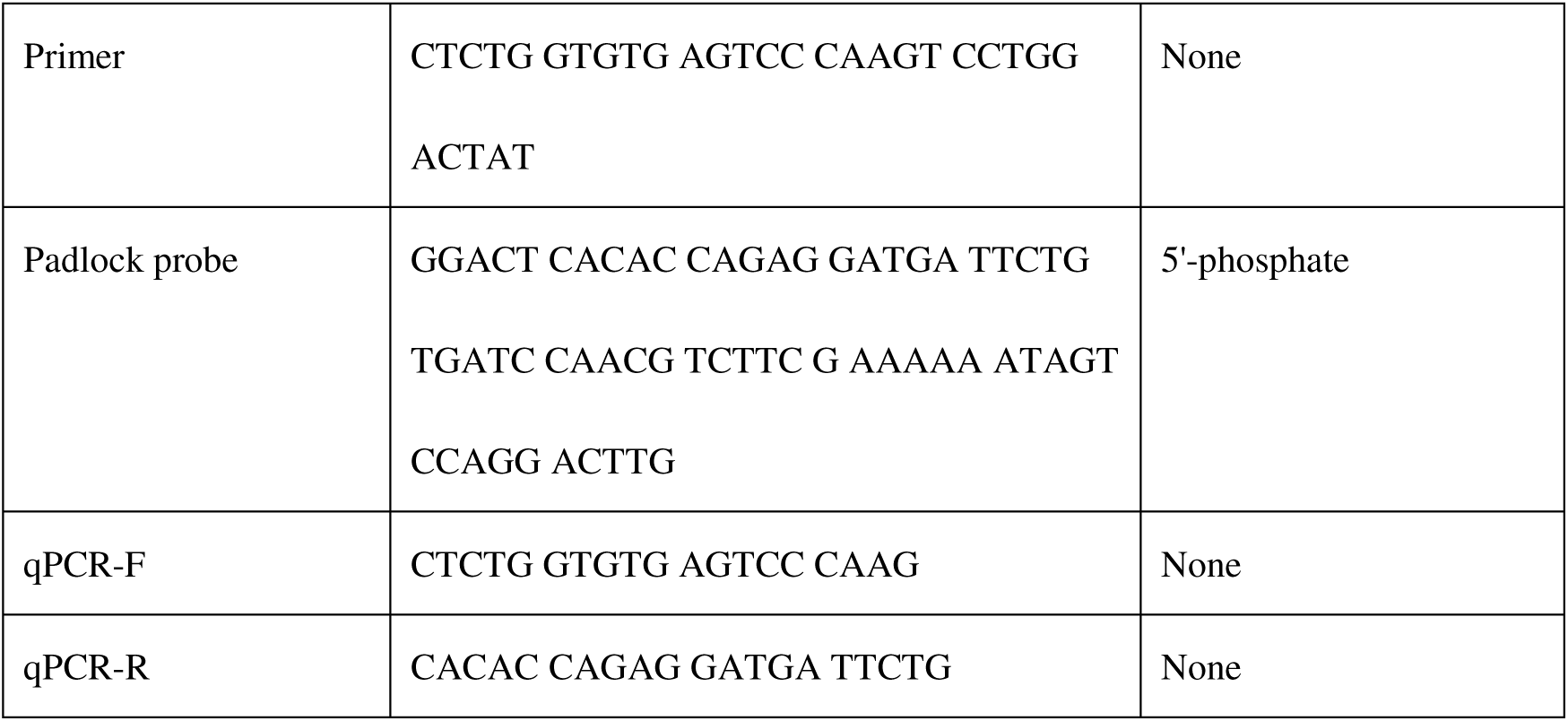

### Mitochondrial DNA copy number quantitation

Mitochondrial DNA (mtDNA) copy number was determined by a quantitative polymerase chain reaction (qPCR) assay with ABI SYBR Green system (Applied Biosystems). 10 ng of total DNA from each cell was used as input for a 10 μL reaction.

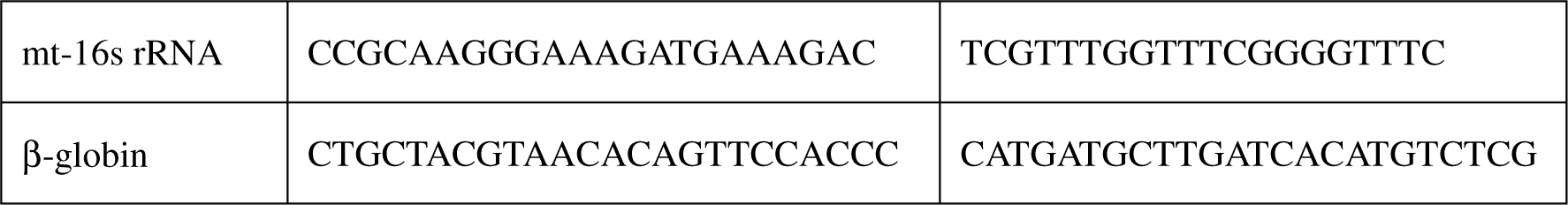

### DNA Fiber Assay

The DNA fiber assay was performed as described previously (Jackson & Pombo, 1998). Briefly, cells were sequentially treated with the medium containing 25 μM of CldU (Sigma-Aldrich, C6891) and then 250 μM of IdU (Sigma-Aldrich, I7125) for 20 min. After washing, cells were suspended in PBS (1000 cells/ μL). 2 μL of cell suspension was spotted on slides. After 5 min, 10 μL spread solution (200 mM Tris-HCl, pH 7.5, 50 mM EDTA, and 0.5% SDS) was spotted on the cell droplet and spread on slides. The slides were then fixed with methanol/acetic acid (3:1), followed by denaturation with 2.5 M HCl. After blocking with 5 % BSA in TBST, slides were stained with rat anti-BrdU antibody (which detects CldU but not IdU, OBT0030, 1:2,000, AbDSerotec) and mouse anti-BrdU antibody (which detects IdU but not CldU, 7580, 1:1,000, BD Biosciences) at 4℃ for overnight. The next day, slides were washed with TBST, followed by staining with the secondary antibodies of goat anti-rat IgG conjugated with TRITC (Jackson Lab) and goat anti-mouse IgG conjugated with FITC. Images of DNA fibers were acquired using a microscope (Olympus). The lengths of red- and green-labeled fibers were determined by FluoView3.0 software (Olympus).

### RNA sequencing analysis

A total amount of 1 μg total RNA per sample was used for sequencing libraries generation by using KAPA mRNA HyperPrep Kit (KAPA Biosystems, Roche, Basel, Switzerland) following manufacturer’s recommendations and index codes were added to attribute sequences to each sample. Briefly, purified mRNA by magnetic oligo-dT was fragmented by high temperature in the presence of Mg^2+^ contained 1x KAPA Fragment, Prime and Elute Buffer. The first strand cDNA was synthesized by using random hexamer priming, followed by 2nd strand synthesis and A-tailing together with dUTP incorporation and adding dAMP at the 3’ ends to form dscDNA. The dsDNA adapter with 3’dTMP overhangs were ligated to dscDNA to generate the library fragments carrying the adapters. In order to select cDNA fragments of preferentially 300∼400bp in length, the library fragments were purified with KAPA Pure Beads system (KAPA Biosystems, Roche, Basel, Switzerland). The library carrying appropriate adapter sequences at both ends was amplified using KAPA HiFi HotStart ReadyMix (KAPA Biosystems, Roche, Basel, Switzerland) along with library amplification primers. The strand marked with dUTP in not amplified, allowing strand-specific sequencing. At last, PCR products were purified using KAPA Pure Beads system and the library quality was assessed on the Qsep 100 DNA/RNA Analyzer (BiOptic Inc., Taiwan). The PCR products were subjected to high-throughput sequencing (Illumina NovaSeq 6000 platform).

The raw sequenced reads were obtained by CASAVA base calling and stored as FASTQ format. After filtered low-quality reads, trim adaptor sequences, and eliminate poor-quality bases by Trimmomatic, the obtained high-quality data (clean reads) was used for subsequent analysis. Read pairs from each sample were aligned to the reference *Mus musculus* genome by the HISAT2 software. FeatureCounts was used to count the reads numbers mapped to individual genes. The quantification and normalization (DESeq method) and further downstream analyses of identification of differentially expressed genes (DEGs) were done by using R package. The resulting p-values were adjusted using the Benjamini and Hochberg’s approach for controlling the FDR. Genes exhibiting false discovery rate (FDR)-adjusted P value < 0.05 and the absolute fold changes (FC) > 2 were considered significant and subjected to further analysis. GO and KEGG pathway enrichment analysis of DEGs were conducted using clusterProfiler. Gene set enrichment analysis (GSEA) was performed with 1,000 permutations to identify enriched biological functions and activated pathways from the molecular signatures database (MSigDB), which including hallmark gene sets, positional gene sets, curated gene sets, motif gene sets, computational gene sets, GO gene sets, oncogenic gene sets, and immunologic gene sets.

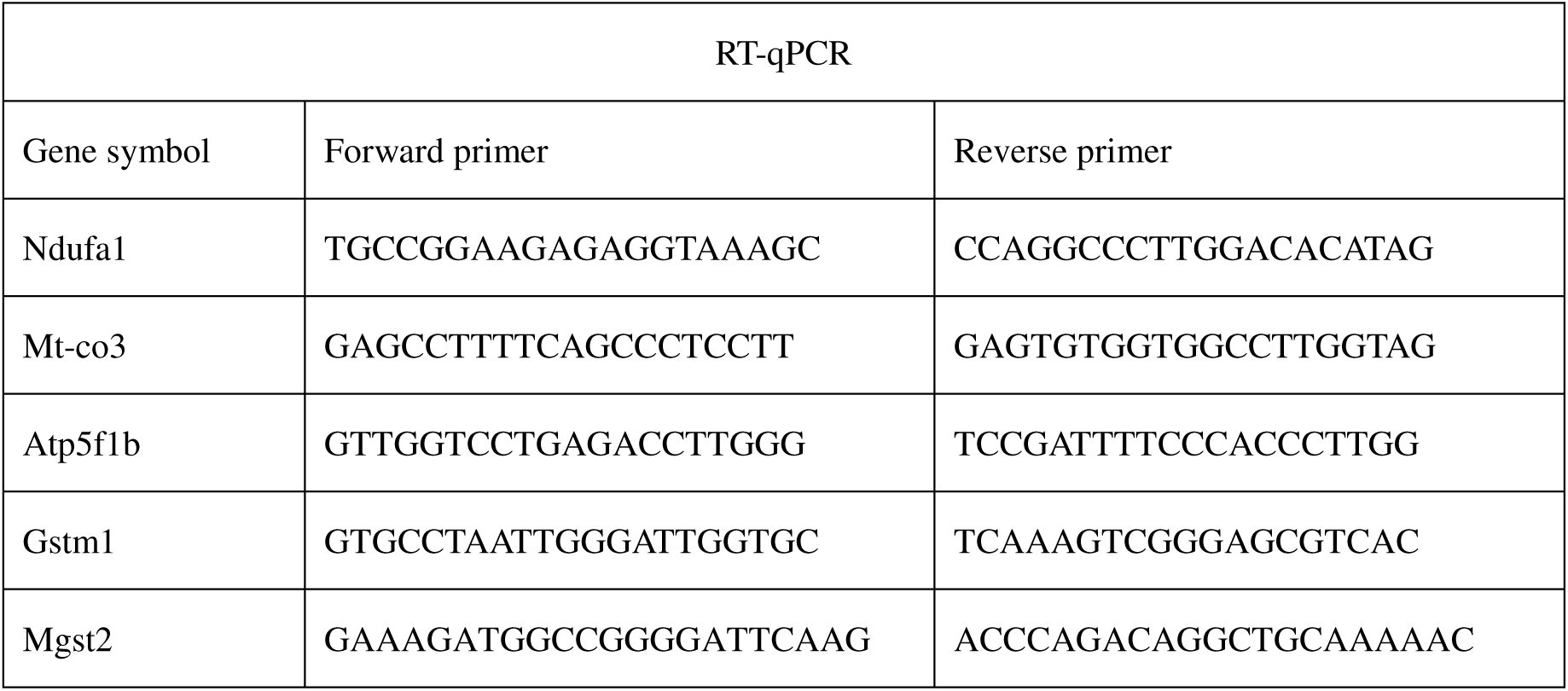
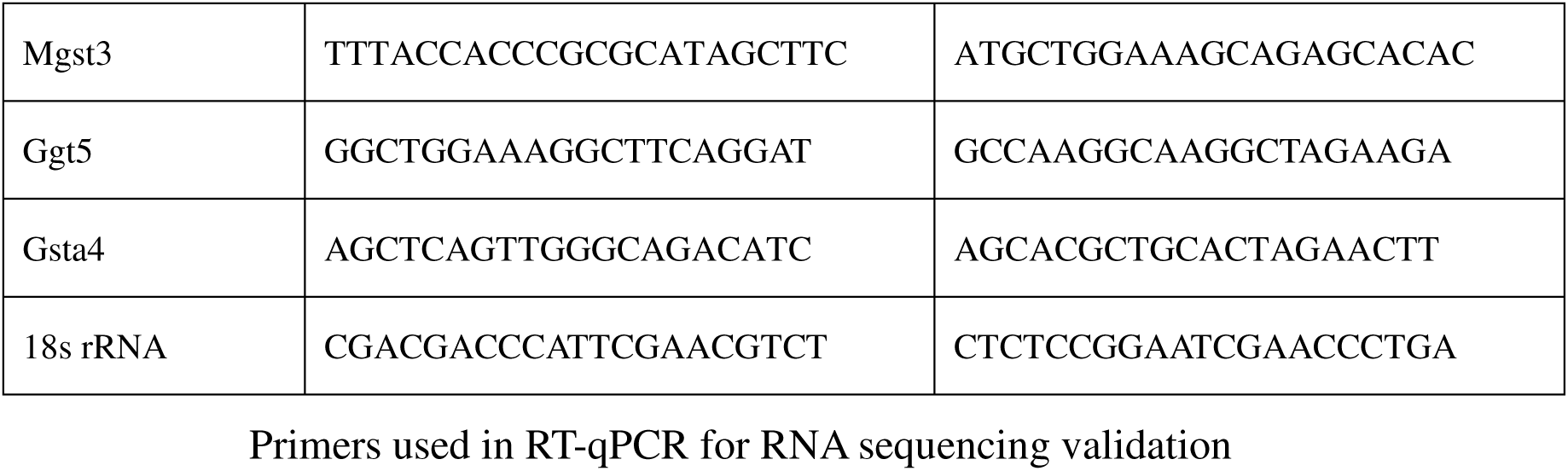

### Statistical analysis

Data are presented as the mean ± SEM Statistical comparison of means was performed using a non-parameter t-test. For mice survival comparison, the Log-rank test was used.

## Acknowledgments

We are grateful for the support from the Laboratory Animal Center at National Taiwan University, College of Medicine and Flow Cytometric Analyzing and Sorting Core of the First Core Laboratory, National Taiwan University College of Medicine for providing the service. We thank Robert Arlinghaus (The University of Texas MD Anderson Cancer Center, Houston, TX, USA) for providing untransformed, WT- and T315I-Bcr-Abl transformed 32D cell lines. We thank Hsueh-Tzu Shih and Chih-Wei Chen for the technical assistance. We thank Gong-Min Lin at Metabolomics Core in Academia Sinica for the assistance of metabolomics analysis and ^13^C metabolite tracing. We also thank Dr. Yu-Lun Kuo at BIOTOOLS Co., Ltd. in Taiwan for kindly supporting analysis RNA sequencing. This research is supported by grants from the Ministry of Science and Technology, Taiwan (MOST 110-2326-B-002-010 and MOST 110–2634- F-002-044).

## Author Contribution

C.Y.H. performed protein purification, enzymatic kinetics, viability, co-factors, metabolic analysis, DNA fiber assay, animal experiments, data analysis, organization and paper writing. S.Y.W. and Y.H.C. performed animal experiments. H.Y.W. performed dNTP quantitation. Z.Y.L. performed metabolomics analysis and ^13^C metabolite tracing experiments. Y.H.C. also performed the Comet assay and DNA fiber analysis. J.M.F and T.J.Y synthesized compounds. C.M.H. tested drugs susceptibility. Z.F.C. conceived, designed, and supervised this project and wrote the paper.

## The Paper explained

### Problem

The acquirement of T315I mutation of Bcr-Abl in chronic myeloid leukemia (CML) leads to resistance to tyrosine kinase inhibitors (TKI). Ponatinib, the inhibitor of T315I- Bcr-Abl, causes severe side-effect. Therefore, developing the therapy for eradicating T315I-Bcr-Abl CML is an unmet need. Metabolism has been suggested as a potential therapeutic target in CML. However, it is unknown what the metabolic vulnerability in T315I-Bcr-Abl cells is.

### Results

We developed a small compound, JMF4073, which inhibited pyrimidylate kinases, to block dNTP synthesis. ROS-induced replication stress in WT-Bcr-Abl-transformed cells causes vulnerability to JMF4073. However, T315I-Bcr-Abl-transformed cells were less vulnerable to JMF4073. We found that WT- and T315I-Bcr-Abl-transformed myeloid cells were different in dNTP pools, replication stress, and Sirt1-regulated mitochondrial respiration. Metabolic and RNA sequencing analyses revealed that T315I mutation caused reductive metabolic shifts to increase glutathione and dNTP levels, making T315I-Bcr-Abl cells less sensitive to JMF4073. However, blocking mitochondrial pyruvate uptake by UK5099 converted T315I-Bcr-Abl cells sensitive to JMF4073 by increasing oxidative load in mitochondria, reducing glutathione and increasing the replication stress. The combination of JMF4073 with UK-5099 depleted dNTPs and showed in vivo eradication of T315I-Bcr-Abl-CML.

### Impact

The impacts of this study are three-folded. First, ROS-induced replication stress causes Bcr-Abl-myeloid cells vulnerable to the inhibitor of pyrimidylate kinases. Second, T315I mutation renders Bcr-Abl-transformed cells deficient of Sirt1-regulated mitochondria and shifted to reductive metabolism without replication stress. Third, cotargeting pyrimidylate kinases and mitochondrial pyruvate uptake induce ROS-dependent replication stress with dNTP depletion in T315I-Bcr-Abl-CML, suggesting a therapeutic option.

## Data availability

All data associated with this study are present in this paper or the supplementary information. RNA sequencing data are deposited in the National Center for Biotechnology Information’s Sequence Read Archive (PRJNA812502) (https://www.ncbi.nlm.nih.gov/sra/?term=PRJNA812502).

## Reference

Abraham A, Qiu S, Chacko BK, Li H, Paterson A, He J, Agarwal P, Shah M, Welner R, Darley-Usmar VM, Bhatia R (2019) SIRT1 regulates metabolism and leukemogenic potential in CML stem cells. J Clin Invest 129: 2685–2701

Altman BJ, Stine ZE, Dang CV (2016) From Krebs to clinic: glutamine metabolism to cancer therapy. Nat Rev Cancer 16: 749

Alvarez-Calderon F, Gregory MA, Pham-Danis C, DeRyckere D, Stevens BM, Zaberezhnyy V, Hill AA, Gemta L, Kumar A, Kumar V, Wempe MF, Pollyea DA, Jordan CT, Serkova NJ, Graham DK, DeGregori J (2015) Tyrosine kinase inhibition in leukemia induces an altered metabolic state sensitive to mitochondrial perturbations. Clin Cancer Res 21: 1360–72

Bajzikova M, Kovarova J, Coelho AR, Boukalova S, Oh S, Rohlenova K, Svec D, Hubackova S, Endaya B, Judasova K, Bezawork-Geleta A, Kluckova K, Chatre L, Zobalova R, Novakova A, Vanova K, Ezrova Z, Maghzal GJ, Magalhaes Novais S, Olsinova M et al. (2019) Reactivation of Dihydroorotate Dehydrogenase-Driven Pyrimidine Biosynthesis Restores Tumor Growth of Respiration-Deficient Cancer Cells. Cell Metab 29: 399–416 e10

Ben-Sahra I, Howell JJ, Asara JM, Manning BD (2013) Stimulation of de novo pyrimidine synthesis by growth signaling through mTOR and S6K1. Science 339: 1323–8

Braun TP, Eide CA, Druker BJ (2020) Response and Resistance to BCR-ABL1- Targeted Therapies. Cancer Cell 37: 530–542

Brown KK, Spinelli JB, Asara JM, Toker A (2017) Adaptive Reprogramming of De Novo Pyrimidine Synthesis Is a Metabolic Vulnerability in Triple-Negative Breast Cancer. Cancer Discov 7: 391–399

Chen CW, Tsao N, Huang LY, Yen Y, Liu X, Lehman C, Wang YH, Tseng MC, Chen YJ, Ho YC, Chen CF, Chang ZF (2016a) The Impact of dUTPase on Ribonucleotide Reductase-Induced Genome Instability in Cancer Cells. Cell Rep 16: 1287–1299

Chen YH, Hsu HY, Yeh MT, Chen CC, Huang CY, Chung YH, Chang ZF, Kuo WC, Chan NL, Weng JH, Chung BC, Chen YJ, Jian CB, Shen CC, Tai HC, Sheu SY, Fang JM (2016b) Chemical Inhibition of Human Thymidylate Kinase and Structural Insights into the Phosphate Binding Loop and Ligand-Induced Degradation. J Med Chem 59: 9906–9918

D’Angiolella V, Donato V, Forrester FM, Jeong YT, Pellacani C, Kudo Y, Saraf A, Florens L, Washburn MP, Pagano M (2012) Cyclin F-mediated degradation of ribonucleotide reductase M2 controls genome integrity and DNA repair. Cell 149: 1023–34

de Beauchamp L, Himonas E, Helgason GV (2022) Mitochondrial metabolism as a potential therapeutic target in myeloid leukaemia. Leukemia 36: 1–12

Deguchi Y, Kimura S, Ashihara E, Niwa T, Hodohara K, Fujiyama Y, Maekawa T (2008) Comparison of imatinib, dasatinib, nilotinib and INNO-406 in imatinib-resistant cell lines. Leuk Res 32: 980–3

Dorsey JF, Cunnick JM, Mane SM, Wu J (2002) Regulation of the Erk2-Elk1 signaling pathway and megakaryocytic differentiation of Bcr-Abl(+) K562 leukemic cells by Gab2. Blood 99: 1388–97

Druker BJ, Talpaz M, Resta DJ, Peng B, Buchdunger E, Ford JM, Lydon NB, Kantarjian H, Capdeville R, Ohno-Jones S, Sawyers CL (2001) Efficacy and safety of a specific inhibitor of the BCR-ABL tyrosine kinase in chronic myeloid leukemia. N Engl J Med 344: 1031–7

Druker BJ, Tamura S, Buchdunger E, Ohno S, Segal GM, Fanning S, Zimmermann J, Lydon NB (1996) Effects of a selective inhibitor of the Abl tyrosine kinase on the growth of Bcr-Abl positive cells. Nat Med 2: 561–6

Flis K, Irvine D, Copland M, Bhatia R, Skorski T (2012) Chronic myeloid leukemia stem cells display alterations in expression of genes involved in oxidative phosphorylation. Leuk Lymphoma 53: 2474–8

Griswold IJ, MacPartlin M, Bumm T, Goss VL, O’Hare T, Lee KA, Corbin AS, Stoffregen EP, Smith C, Johnson K, Moseson EM, Wood LJ, Polakiewicz RD, Druker BJ, Deininger MW (2006) Kinase domain mutants of Bcr-Abl exhibit altered transformation potency, kinase activity, and substrate utilization, irrespective of sensitivity to imatinib. Mol Cell Biol 26: 6082–93

Halazonetis TD, Gorgoulis VG, Bartek J (2008) An oncogene-induced DNA damage model for cancer development. Science 319: 1352–5

Hatzivassiliou G, Zhao F, Bauer DE, Andreadis C, Shaw AN, Dhanak D, Hingorani SR, Tuveson DA, Thompson CB (2005) ATP citrate lyase inhibition can suppress tumor cell growth. Cancer Cell 8: 311–21

Herzig S, Shaw RJ (2018) AMPK: guardian of metabolism and mitochondrial homeostasis. Nat Rev Mol Cell Biol 19: 121–135

Hills SA, Diffley JF (2014) DNA replication and oncogene-induced replicative stress. Curr Biol 24: R435–44

Hu CM, Tsao N, Wang YT, Chen YJ, Chang ZF (2019) Thymidylate kinase is critical for DNA repair via ATM-dependent Tip60 complex formation. FASEB J 33: 2017–2025

Hu CM, Yeh MT, Tsao N, Chen CW, Gao QZ, Chang CY, Lee MH, Fang JM, Sheu SY, Lin CJ, Tseng MC, Chen YJ, Chang ZF (2012a) Tumor cells require thymidylate kinase to prevent dUTP incorporation during DNA repair. Cancer Cell 22: 36–50

Hu Y, Lu W, Chen G, Wang P, Chen Z, Zhou Y, Ogasawara M, Trachootham D, Feng L, Pelicano H, Chiao PJ, Keating MJ, Garcia-Manero G, Huang P (2012b) K-ras(G12V) transformation leads to mitochondrial dysfunction and a metabolic switch from oxidative phosphorylation to glycolysis. Cell Res 22: 399–412

Huang Y, Lu J, Zhan L, Wang M, Shi R, Yuan X, Gao X, Liu X, Zang J, Liu W, Yao X (2021) Resveratrol-induced Sirt1 phosphorylation by LKB1 mediates mitochondrial metabolism. J Biol Chem 297: 100929

Jackson DA, Pombo A (1998) Replicon clusters are stable units of chromosome structure: evidence that nuclear organization contributes to the efficient activation and propagation of S phase in human cells. J Cell Biol 140: 1285–95

Ke PY, Kuo YY, Hu CM, Chang ZF (2005) Control of dTTP pool size by anaphase promoting complex/cyclosome is essential for the maintenance of genetic stability. Genes Dev 19: 1920–33

Kovacevic Z (1971) The pathway of glutamine and glutamate oxidation in isolated mitochondria from mammalian cells. Biochem J 125: 757–63

Kuntz EM, Baquero P, Michie AM, Dunn K, Tardito S, Holyoake TL, Helgason GV, Gottlieb E (2017) Targeting mitochondrial oxidative phosphorylation eradicates therapy-resistant chronic myeloid leukemia stem cells. Nat Med 23: 1234–1240

Lagouge M, Argmann C, Gerhart-Hines Z, Meziane H, Lerin C, Daussin F, Messadeq N, Milne J, Lambert P, Elliott P, Geny B, Laakso M, Puigserver P, Auwerx J (2006) Resveratrol improves mitochondrial function and protects against metabolic disease by activating SIRT1 and PGC-1alpha. Cell 127: 1109–22

Lan F, Cacicedo JM, Ruderman N, Ido Y (2008) SIRT1 modulation of the acetylation status, cytosolic localization, and activity of LKB1. Possible role in AMP-activated protein kinase activation. J Biol Chem 283: 27628–27635

Le TM, Poddar S, Capri JR, Abt ER, Kim W, Wei L, Uong NT, Cheng CM, Braas D, Nikanjam M, Rix P, Merkurjev D, Zaretsky J, Kornblum HI, Ribas A, Herschman HR, Whitelegge J, Faull KF, Donahue TR, Czernin J et al. (2017) ATR inhibition facilitates targeting of leukemia dependence on convergent nucleotide biosynthetic pathways. Nat Commun 8: 241

Lecona E, Fernandez-Capetillo O (2018) Targeting ATR in cancer. Nat Rev Cancer 18: 586–595

Leong D, Aghel N, Hillis C, Siegal D, Karampatos S, Rangarajan S, Pond G, Seow H (2021) Tyrosine kinase inhibitors in chronic myeloid leukaemia and emergent cardiovascular disease. Heart 107: 667–673

Li L, Wang L, Li L, Wang Z, Ho Y, McDonald T, Holyoake TL, Chen W, Bhatia R (2012) Activation of p53 by SIRT1 inhibition enhances elimination of CML leukemia stem cells in combination with imatinib. Cancer Cell 21: 266–81

Locasale JW (2013) Serine, glycine and one-carbon units: cancer metabolism in full circle. Nat Rev Cancer 13: 572–83

Mian AA, Schull M, Zhao Z, Oancea C, Hundertmark A, Beissert T, Ottmann OG, Ruthardt M (2009) The gatekeeper mutation T315I confers resistance against small molecules by increasing or restoring the ABL-kinase activity accompanied by aberrant transphosphorylation of endogenous BCR, even in loss-of-function mutants of BCR/ABL. Leukemia 23: 1614–21

Nieborowska-Skorska M, Kopinski PK, Ray R, Hoser G, Ngaba D, Flis S, Cramer K, Reddy MM, Koptyra M, Penserga T, Glodkowska-Mrowka E, Bolton E, Holyoake TL, Eaves CJ, Cerny-Reiterer S, Valent P, Hochhaus A, Hughes TP, van der Kuip H, Sattler M et al. (2012) Rac2-MRC-cIII-generated ROS cause genomic instability in chronic myeloid leukemia stem cells and primitive progenitors. Blood 119: 4253–63

Niida H, Katsuno Y, Sengoku M, Shimada M, Yukawa M, Ikura M, Ikura T, Kohno K, Shima H, Suzuki H, Tashiro S, Nakanishi M (2010) Essential role of Tip60-dependent recruitment of ribonucleotide reductase at DNA damage sites in DNA repair during G1 phase. Genes Dev 24: 333–8

Nowicki MO, Falinski R, Koptyra M, Slupianek A, Stoklosa T, Gloc E, Nieborowska-Skorska M, Blasiak J, Skorski T (2004) BCR/ABL oncogenic kinase promotes unfaithful repair of the reactive oxygen species-dependent DNA double-strand breaks. Blood 104: 3746–53

Ren R (2005) Mechanisms of BCR-ABL in the pathogenesis of chronic myelogenous leukaemia. Nat Rev Cancer 5: 172–83

Robitaille AM, Christen S, Shimobayashi M, Cornu M, Fava LL, Moes S, Prescianotto-Baschong C, Sauer U, Jenoe P, Hall MN (2013) Quantitative phosphoproteomics reveal mTORC1 activates de novo pyrimidine synthesis. Science 339: 1320–3

Rodgers JT, Lerin C, Haas W, Gygi SP, Spiegelman BM, Puigserver P (2005) Nutrient control of glucose homeostasis through a complex of PGC-1alpha and SIRT1. Nature 434: 113–8

Rosti G, Castagnetti F, Gugliotta G, Baccarani M (2017) Tyrosine kinase inhibitors in chronic myeloid leukaemia: which, when, for whom? Nat Rev Clin Oncol 14: 141–154

Rowley JD (1973) Letter: A new consistent chromosomal abnormality in chronic myelogenous leukaemia identified by quinacrine fluorescence and Giemsa staining. Nature 243: 290–3

Ruderman NB, Xu XJ, Nelson L, Cacicedo JM, Saha AK, Lan F, Ido Y (2010) AMPK and SIRT1: a long-standing partnership? Am J Physiol Endocrinol Metab 298: E751–60

Sabharwal SS, Schumacker PT (2014) Mitochondrial ROS in cancer: initiators, amplifiers or an Achilles’ heel? Nat Rev Cancer 14: 709–21

Salesse S, Verfaillie CM (2002) BCR/ABL: from molecular mechanisms of leukemia induction to treatment of chronic myelogenous leukemia. Oncogene 21: 8547–59

Sattler M, Mohi MG, Pride YB, Quinnan LR, Malouf NA, Podar K, Gesbert F, Iwasaki H, Li S, Van Etten RA, Gu H, Griffin JD, Neel BG (2002) Critical role for Gab2 in transformation by BCR/ABL. Cancer Cell 1: 479–92

Sawyers CL (1999) Chronic myeloid leukemia. N Engl J Med 340: 1330–40

Segura-Pena D, Sekulic N, Ort S, Konrad M, Lavie A (2004) Substrate-induced conformational changes in human UMP/CMP kinase. J Biol Chem 279: 33882–9

Skaggs BJ, Gorre ME, Ryvkin A, Burgess MR, Xie Y, Han Y, Komisopoulou E, Brown LM, Loo JA, Landaw EM, Sawyers CL, Graeber TG (2006) Phosphorylation of the ATP-binding loop directs oncogenicity of drug-resistant BCR-ABL mutants. Proc Natl Acad Sci U S A 103: 19466–71

Smith B, Schafer XL, Ambeskovic A, Spencer CM, Land H, Munger J (2016) Addiction to Coupling of the Warburg Effect with Glutamine Catabolism in Cancer Cells. Cell Rep 17: 821–836

Tang BL (2016) Sirt1 and the Mitochondria. Mol Cells 39: 87–95

Tompkins SC, Sheldon RD, Rauckhorst AJ, Noterman MF, Solst SR, Buchanan JL, Mapuskar KA, Pewa AD, Gray LR, Oonthonpan L, Sharma A, Scerbo DA, Dupuy AJ, Spitz DR, Taylor EB (2019) Disrupting Mitochondrial Pyruvate Uptake Directs Glutamine into the TCA Cycle away from Glutathione Synthesis and Impairs Hepatocellular Tumorigenesis. Cell Rep 28: 2608–2619 e6

Vander Heiden MG, Cantley LC, Thompson CB (2009) Understanding the Warburg effect: the metabolic requirements of cell proliferation. Science 324: 1029–33

Vertessy BG, Toth J (2009) Keeping uracil out of DNA: physiological role, structure and catalytic mechanism of dUTPases. Acc Chem Res 42: 97–106

Wang HY, Hsin P, Huang CY, Chang ZF (2021) A Convenient and Sensitive Method for Deoxynucleoside Triphosphate Quantification by the Combination of Rolling Circle Amplification and Quantitative Polymerase Chain Reaction. Anal Chem 93: 14247–14255

Ward PS, Thompson CB (2012) Metabolic reprogramming: a cancer hallmark even warburg did not anticipate. Cancer Cell 21: 297–308

Xiao B, Sanders MJ, Carmena D, Bright NJ, Haire LF, Underwood E, Patel BR, Heath RB, Walker PA, Hallen S, Giordanetto F, Martin SR, Carling D, Gamblin SJ (2013) Structural basis of AMPK regulation by small molecule activators. Nat Commun 4: 3017

Xiao B, Sanders MJ, Underwood E, Heath R, Mayer FV, Carmena D, Jing C, Walker PA, Eccleston JF, Haire LF, Saiu P, Howell SA, Aasland R, Martin SR, Carling D, Gamblin SJ (2011) Structure of mammalian AMPK and its regulation by ADP. Nature 472: 230–3

Xie S, Wang Y, Liu J, Sun T, Wilson MB, Smithgall TE, Arlinghaus RB (2001) Involvement of Jak2 tyrosine phosphorylation in Bcr-Abl transformation. Oncogene 20: 6188–95

Yang M, Vousden KH (2016) Serine and one-carbon metabolism in cancer. Nat Rev Cancer 16: 650–62

Yuan H, Wang Z, Li L, Zhang H, Modi H, Horne D, Bhatia R, Chen W (2012) Activation of stress response gene SIRT1 by BCR-ABL promotes leukemogenesis. Blood 119: 1904–14

Zhang W, Tan S, Paintsil E, Dutschman GE, Gullen EA, Chu E, Cheng YC (2011) Analysis of deoxyribonucleotide pools in human cancer cell lines using a liquid chromatography coupled with tandem mass spectrometry technique. Biochem Pharmacol 82: 411–7

